# *Aedes aegypti VLG-1* challenges the assumed antiviral nature of *Vago* genes

**DOI:** 10.1101/2024.07.01.601473

**Authors:** Elodie Couderc, Anna B. Crist, Josquin Daron, Hugo Varet, Femke A. H. van Hout, Pascal Miesen, Umberto Palatini, Stéphanie Dabo, Thomas Vial, Louis Lambrechts, Sarah H. Merkling

**Affiliations:** Institut Pasteur, Université Paris Cité, CNRS UMR2000, Insect-Virus Interactions Unit, 75015 Paris, France; Sorbonne Université, Collège Doctoral, 75005 Paris, France; Institut Pasteur, Université Paris Cité, Bioinformatics and Biostatistics Hub, 75015 Paris, France; Department of Medical Microbiology, Radboud University Medical Center, P.O. box 9101 6500 HB Nijmegen, The Netherlands; Laboratory of Neurogenetics and Behavior, The Rockefeller University, New York, NY 10065, USA

## Abstract

Arthropod-borne viruses (arboviruses) such as dengue virus (DENV) and Zika virus (ZIKV) pose a significant threat to global health. Novel approaches to control the spread of arboviruses focus on harnessing the antiviral immune system of their primary vector, the *Aedes aegypti* mosquito. In arthropods, genes of the *Vago* family are often presented as analogs of mammalian cytokines with potential antiviral functions, but the role of *Vago* genes upon virus infection in *Ae. aegypti* is largely unknown. We conducted a phylogenetic analysis of the *Vago* gene family in Diptera, which led us to focus on a *Vago*-like gene that we named *VLG-1*. Using CRISPR/Cas9-mediated gene editing, we generated a *VLG-1* mutant line of *Ae. aegypti*, which revealed a broad impact of *VLG-1* on the mosquito transcriptome, affecting several biological processes potentially related to viral replication, including the oxidative stress response. Surprisingly, experimental viral challenge of the *VLG-1* mutant line indicated a modest proviral role for this gene during DENV and ZIKV infections *in vivo*. In the absence of *VLG-1*, virus dissemination throughout the mosquito’s body was slightly impaired, albeit not altering virus transmission rates. Our results challenge the conventional understanding of *Vago*-like genes as antiviral factors and underscore the need for further *in vivo* research to elucidate the molecular mechanisms underlying mosquito-arbovirus interactions.

## INTRODUCTION

Arthropod-borne viruses (arboviruses) pose a significant threat to global health, causing numerous human diseases with substantial morbidity and mortality. Among the most medically significant arboviruses are the mosquito-borne flaviviruses [1]. For instance, dengue virus (DENV) infects approximately 400 million people each year and is responsible for about 100 million symptomatic cases [2-4]. In addition, Zika virus (ZIKV) emerged in more than 87 countries and territories in the last 15 years, causing severe neuropathologies and birth defects [5]. DENV and ZIKV are primarily transmitted by *Aedes aegypti* (*Ae. aegypti*), a mosquito species found throughout the tropics and subtropics whose range is expected to further expand with global change [6, 7]. To date, there are no globally approved vaccines or specific antivirals against these diseases. Traditional vector control methods are limited in efficacy because of the emergence of insecticide-resistant mosquitoes. Thus, the release of lab-modified mosquitoes that are incapable of transmitting viruses is an alternative strategy for reducing the incidence of human arboviral diseases [8, 9]. The development of such novel interventions is conditioned by the identification of optimal target genes that mediate interactions between mosquitoes and viruses [9, 10].

Female mosquitoes acquire arboviruses by biting and blood feeding on viremic vertebrate hosts. The bloodmeal is digested in the midgut, where viral particles infect epithelial cells [11, 12]. The virus then disseminates through the mosquito body, likely via circulating immune cells called hemocytes [13-15], until it reaches the salivary glands, where it replicates before being released in the saliva [16]. The mosquito can transmit the virus to the next host during a subsequent blood-feeding event [13]. Within mosquitoes, virus infection and dissemination are hindered by physical tissue barriers [13] and innate immune pathways, including RNA interference (RNAi) [17-19], Janus kinase/signal transducers and activators of transcription (JAK-STAT), Toll, and immune deficiency (IMD) pathways, which are activated upon viral detection and trigger the production of effector molecules that can inhibit viral replication [20-22]. Most of our knowledge about antiviral immunity in mosquitoes is derived from pioneering work in the model organism *Drosophila melanogaster*. However, fruit flies are neither arbovirus vectors, nor hematophagous insects, leaving our understanding of mosquito antiviral responses incomplete [10, 20, 23].

For instance, only a few studies have investigated the role of immunoregulatory genes with cytokine-like functions, such as *Vago* genes, in mosquito immunity. The first *Vago* gene was identified in *D. melanogaster* [24] and encodes a secreted antiviral protein induced upon infection by *Drosophila* C virus (DCV) [25]. In *D. melanogaster*, *Vago* induction in response to DCV infection requires the RNAi gene *Dicer2* [25]. In mosquitoes, a *Vago* gene called *CxVago,* was shown to limit viral replication in *Culex* mosquito cells infected with the flavivirus West Nile virus (WNV) [26]. In addition, WNV infection was found to induce the expression of *CxVago* in a *Dicer2*-dependent manner, leading to secretion of the protein and activation of the JAK-STAT pathway via an unknown non-canonical receptor [26]. *Rel2* and *TRAF* genes were also involved in *CxVago* induction, suggesting a link between *CxVago* induction and NF-κB pathways [27]. However, the antiviral function of *Vago* genes in *Culex* mosquitoes was not investigated *in vivo*. Finally, another study using an *Aedes*-derived cell line reported an antiviral role for a *Vago* gene called *AaeVago1*, in the context of DENV and *Wolbachia* co-infection [28].

The Vago protein family is often referred to as “arthropod cytokines” because they are functionally analogous to mammalian cytokines [26, 29-31]. In dipteran insects (flies and mosquitoes), Vago proteins consist of 100-200 amino acids with a secretion signal peptide and a single domain von Willebrand factor type C (SVWC) functional domain. SVWC proteins, characterized by a repetitive pattern of eight cysteines, represent a broadly conserved protein family in arthropods, associated with responses to environmental challenges, including nutritional stress and microbial infections [29]. Despite their characteristic structural features, the functions of Vago proteins in insects remain elusive, particularly *in vivo*.

Here, we investigated the role of *Vago* genes in *Ae. aegypti* mosquitoes *in vivo* in the context of flavivirus infection. We generated and characterized a mosquito mutant line for the gene that had hitherto been called *AaeVago1* and determined its impact on the mosquito transcriptome. We also investigated its role in infection, systemic dissemination, and transmission of DENV and ZIKV. Unexpectedly, we found a subtle proviral effect of this gene, challenging the hypothesis that genes belonging to the *Vago* family exert exclusively antiviral functions in arthropods.

## RESULTS

### *VLG-1* is a *Vago*-like gene exclusively found in the Culicinae

To investigate the role of *Vago* genes in *Ae. aegypti*, we first reconstituted their evolutionary history (Figure 1A). We identified the homologs of *AAEL000200* and *AAEL000165*, two genes that were previously described as *AaeVago1* and *AaeVago2*, in a panel of Diptera species from the Culicidae family (mosquitoes) and from the *Drosophila* genus, and we determined their phylogenetic relationships at the protein level (Supplementary Figure S1). First, we discovered that the first *Vago* gene characterized in *Drosophila melanogaster* (*DmVago*, *CG2081*) [25] is not the most likely homolog of *AAEL000200* and *AAEL000165.* These two *Ae. aegypti* genes encode proteins that are ∼40-50 amino acid shorter and only share 27% and 24% identity with DmVago, respectively (Figure 1B). Reciprocally, *DmVago* does not have a homolog in the Culicidae sharing at least 30% protein sequence identity. We found that the most likely homolog of *AAEL000200* and *AAEL000165* in the *D. melanogaster* genome is an uncharacterized gene (*CG14132*), which we named “*D. melanogaster Vago-like gene*” (*DmVLG*). *DmVLG* shares 36% and 31% protein identity with *AAEL000200* and *AAEL000165*, respectively (Figure 1B). Thus, we renamed *AAEL000200* “*Ae. aegypti Vago-like gene 1”* (*AaeVLG-1*, referred to later in this study as *VLG-1*) and *AAEL000165* “*Ae. aegypti Vago-like gene 2”* (*AaeVLG-2*, referred to later in this study as *VLG-2*). A summary of our proposed updated designation of *Vago* and *Vago*-like genes is provided in Supplementary Table 1.

**Figure 1.**
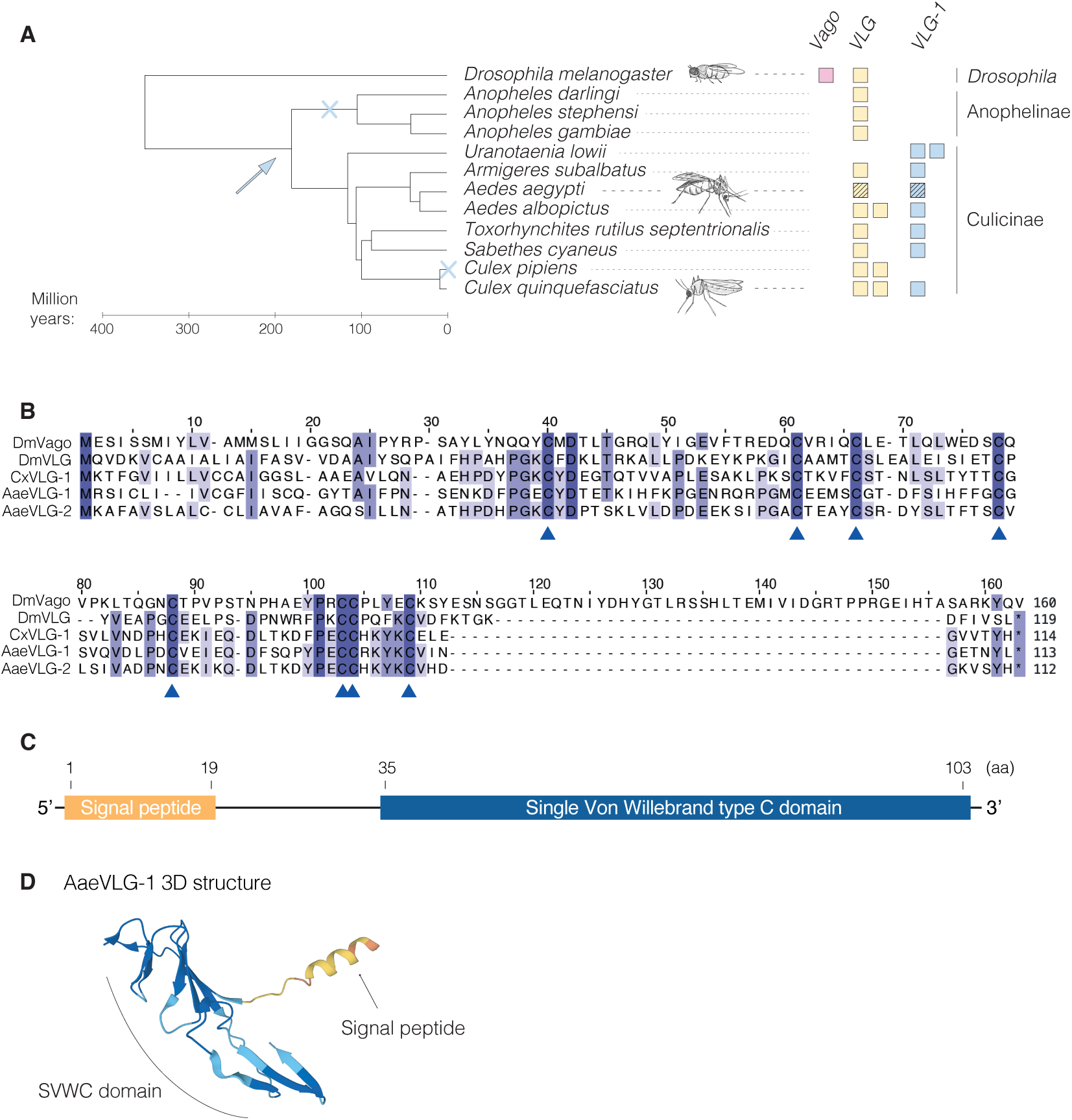
*VLG-1* is a *Vago*-like gene specific to the Culicinae subfamily. (A) Schematic cladogram of the evolutionary history of *Vago*-like gene homologs in Culicidae and *Drosophila* species. The putative origin of duplication of *VLG-1* from the ancestral *VLG,* inferred from the phylogenetic analysis of *Vago*-like gene homologs (Supplementary Figure S1), is indicated with a blue arrow, whereas putative losses of *VLG-1* are indicated with blue crosses. *AaeVLG-2* and *AaeVLG-1* are represented with black-striped yellow and blue squares, respectively. (B) Amino-acid sequence alignment of *Ae. aegypti* VLG-1 (AaeVLG-1, XP 001658930.1) and VLG-2 (isoform RB, XP 001658929.1) proteins with *D. melanogaster* Vago (DmVago, NP_001285106.1), *D. melanogaster* VLG (DmVLG, NP_001097586) and *Culex quinquefasciatus* VLG-1 (XP_001842264). The percentage of identity shared between all sequences for each amino-acid position is represented by shades of colors, ranging from light purple (when 3 out of 5 sequences are identical) to dark purple (when all 5 sequences are identical). The conserved cysteine residues typical of SVWC domains are indicated by blue arrows. (C) Functional domains of AaeVLG-1 with amino-acid (aa) positions. (D) Predicted 3D structure of AaeVLG-1 protein obtained with Alphafold (https://alphafold.ebi.ac.uk/entry/Q17PX2) [77, 78].

The overall topology of the phylogenetic tree of *Vago*-like gene homologs revealed two distinct sister clades among the Culicidae (Supplementary Figure S1). One clade encompasses members of both the Culicinae and Anophelinae subfamilies, including *AaeVLG-2*. The other clade exclusively consists of Culicinae members, including *AaeVLG-1*. The *VLG* clade that includes *AaeVLG-2* likely represents the orthologous group of *DmVLG*, whereas the clade that includes *AaeVLG-1* likely corresponds to *Vago*-like paralogs that arose by duplication of the ancestral *VLG*. This scenario is further supported by the nested and inverted position of the *AaeVLG-1* locus within an intron of *AaeVLG-2*. Our analysis suggests that the duplication occurred prior to the divergence of Anophelinae and Culicinae and was followed by a loss of the duplicated copy in Anophelinae prior to their diversification (Figure 1A and Supplementary Figure S1). Together, our analysis identified *AaeVLG-2* (previously named *AaeVago2* [28]) as the direct ortholog of *DmVLG* in *Ae. aegypti*, and *AaeVLG-1* (previously named *AaeVago1* [28]) as the duplicated copy. Accordingly, we also propose to rename the *Culex quinquefasciatus* gene *CQUJHB003889*, previously known as *CxVago* [26, 27], as *CxVLG-1* because it belongs to the *VLG-1* clade (Supplementary Figure S1 and Supplementary Table 1).

To determine whether the two *Vago*-like copies in the Culicidae family evolved under a different selection regime after the duplication event, we estimated the evolutionary rates of *AaeVLG-1* and *AaeVLG-2*. We computed the ratio of non-synonymous to synonymous substitutions (ω) for all *VLG* homologs of our panel of Culicidae and *Drosophila* species. The ω ratio, also known as dN/dS, indicates the mode and strength of natural selection, where ω=0 means that the gene is under purifying selection, ω=1 indicates neutral selection, and ω>1 indicates diversifying selection. We used a branch model that evaluates the variation of ω within the tree and tests for differences in selection regimes between lineages. According to this model, both *VLG-1* and Culicinae *VLG* are under purifying selection (ω=0.18 and ω=0.15, respectively), but slightly weaker purifying selection than Anophelinae *VLG* (ω=0.1) and *Drosophila VLG* (ω=0.09) (Supplementary Figure S1 and Supplementary Table 3). This analysis suggests that the *VLG* duplication in the Culicinae was followed by relaxed selective pressure on both copies.

At the amino-acid level, *AaeVLG-1* shares 57% identity with *CxVLG-1*, whereas *AaeVLG-2* shares 38% identity with *CxVLG-1* (Figure 1B). *AaeVLG-1* is transcribed into a 451-bp mRNA transcript encoding a protein of 113 amino acids, including a signal peptide, theoretically responsible for addressing the protein to the membrane prior to its secretion, and an SVWC domain with the characteristic eight-cysteine repeat (Figure 1B-D).

### *VLG-1* is persistently induced by bloodmeal ingestion in *Ae. aegypti*

In arthropods, *Vago* genes have been described as factors induced by biotic or abiotic stress [25-29, 32-35]. In *Ae. aegypti*, the potential role of *VLG-1* and *VLG-2* during viral infection has only been investigated *in vitro*, in an *Aedes*-derived cell line [28]. To test whether *VLG-1* and *VLG-2* are induced upon viral infection *in vivo* in *Ae. aegypti*, we exposed mosquitoes to a bloodmeal containing DENV (DENV-1) or a control mock bloodmeal. We quantified the expression of *VLG-1* and *VLG-2* by quantitative RT-PCR (RT-qPCR) in individual midguts, heads, and carcasses (*i.e.,* bodies without midgut and head) at several timepoints post bloodmeal, from day 0 to day 9 (Figure 2). As reported previously [36], we found that in *Ae. aegypti*, overall transcript abundance was ∼2- to 10-fold higher for *VLG-1* than for *VLG-2* across tissues (Figure 2A-F). A mock bloodmeal triggered a persistent up-regulation of *VLG-1* transcription lasting up to 9 days post bloodmeal in carcasses and heads (Figure 2B and 2C). DENV exposure triggered a transient and modest increase in *VLG-1* expression in heads day 2 post bloodmeal (Figure 2C). No differences in *VLG-2* expression levels were detected between the mock and the infectious bloodmeals (Figure 2D-F). Therefore, because *AaeVLG-1* displays higher expression levels than *AaeVLG-2* and is persistently induced by bloodmeal ingestion, we chose to focus our study on the role of *AaeVLG-1* upon virus infection.

**Figure 2.**
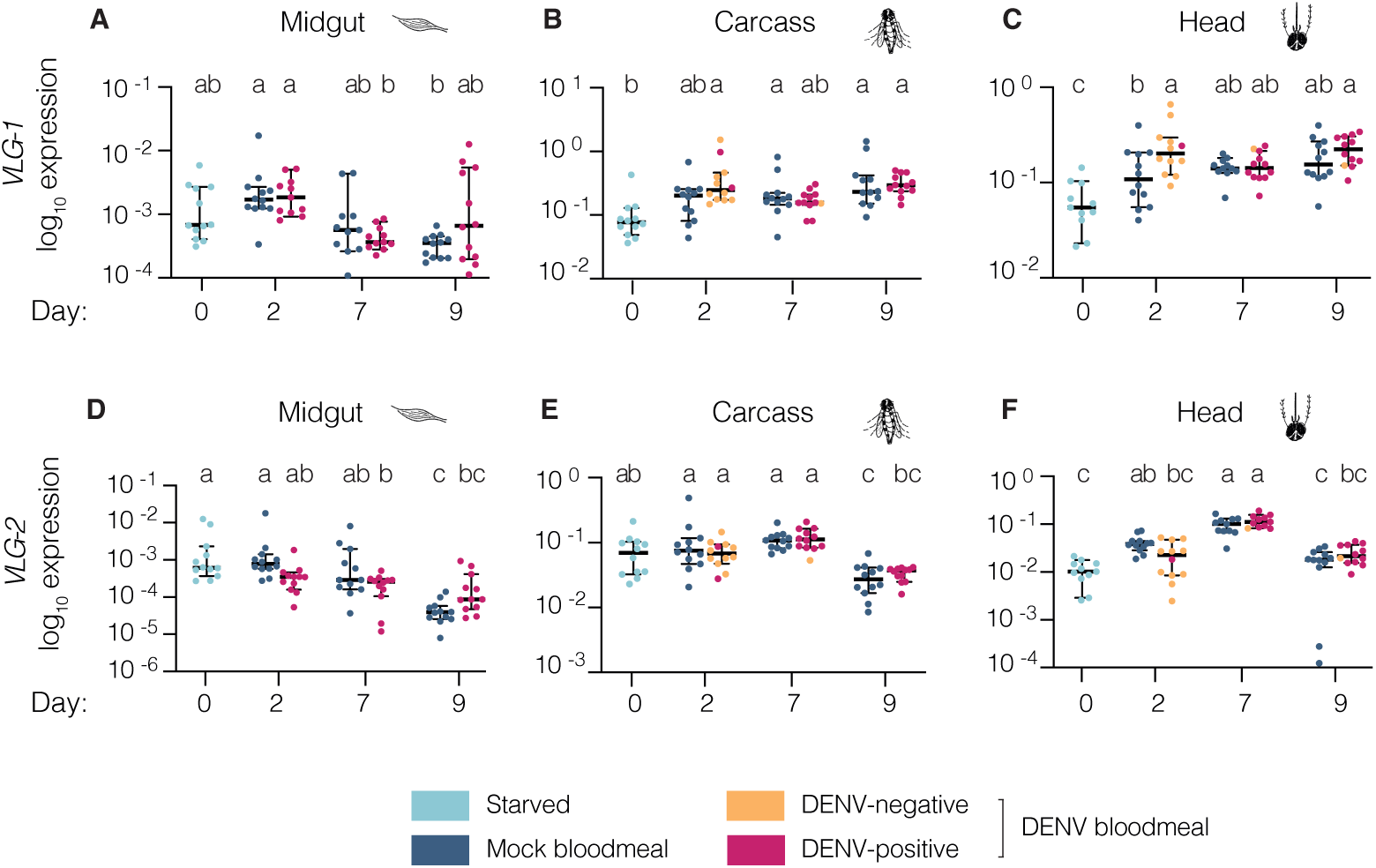
*VLG-1* is persistently induced by bloodmeal ingestion and DENV exposure in non-midgut tissues of *Ae. aegypti*. Expression levels of *AaeVLG-1* (A-C) and *AaeVLG-2* (D-F) were quantified by RT-qPCR in midguts (A,D), carcasses (B,E), and heads (C,F) on 0, 2, 7, and 9 days after ingestion of a mock or DENV-1 infectious bloodmeal, in mosquitoes of the wild-type control line. Tissues from DENV-exposed mosquitoes were sorted into DENV-positive and DENV-negative samples. Gene expression levels are normalized to the ribosomal protein S 17 housekeeping gene (*RPS17*), and expressed as 2^-dCt^, where dCt = Ct*_Gene_* – Ct*_RPS17_*. Each dot represents an individual tissue. Horizontal bars represent medians and vertical bars represent 95% confidence intervals. Statistical significance was determined by a one-way ANOVA after log_10_- transformation of the 2^-dCt^ values, followed by Tukey-Kramer’s HSD test. Statistical significance is represented above the graph using letters; groups that do not share a letter are significantly different.

### *Ae. aegypti VLG-1*^Δ^ mutant mosquitoes do not exhibit major fitness defects

To further investigate the role of *VLG-1* in *Ae. aegypti*, we generated a mutant line by CRISPR/Cas9-mediated gene editing. Shortly, mosquito embryos were microinjected with Cas9 coupled to 3 single-guide RNAs (sgRNAs) targeting 3 *VLG-1* exons together with a repair template (Figure 3A). We isolated one generation zero (G_0_) female carrying a 212-bp (55 amino acids) deletion in the *VLG-1* locus, resulting from a combined 246-bp deletion and a 34-bp insertion from the repair template. This G_0_ female was crossed to wild-type males and the resulting G_1_ males carrying the deletion were crossed to wild-type females for three more generations. G_4_ adults carrying the *VLG-1* mutation at the homozygous state were used to establish a *VLG-1* mutant line that we called *VLG-1*^Δ^. Within the same crossing scheme, we generated a control “sister” line carrying the wild-type version of *VLG-1*. The *VLG-1*^Δ^ line encodes a *VLG-1* protein with only 58 of the 113 original amino acids left and 81% of the SVWC functional domain truncated, suggesting a *VLG-1* loss of function. We found a strong decrease of *VLG-1* transcript abundance in the mutant line relative to the control line, both by RT-qPCR and by RNA sequencing (RNA-seq) (Supplementary Figure S2A-B), which is a hallmark of nonsense-mediated decay of the aberrant mRNA [37]. We also confirmed the absence of detectable VLG-1 protein in the mutant line by Western blot using a previously developed anti-VLG-1 antibody [26] (Supplementary Figure S2C). No off-target effect on *AaeVLG-2* expression was detected by RT-qPCR or RNA-seq in the *VLG-1*^Δ^ mutant line (Supplementary Figure S2D-E). Together, these results strongly suggest that we generated a *bona fide* knock-out *VLG-1* mutant line.

**Figure 3.**
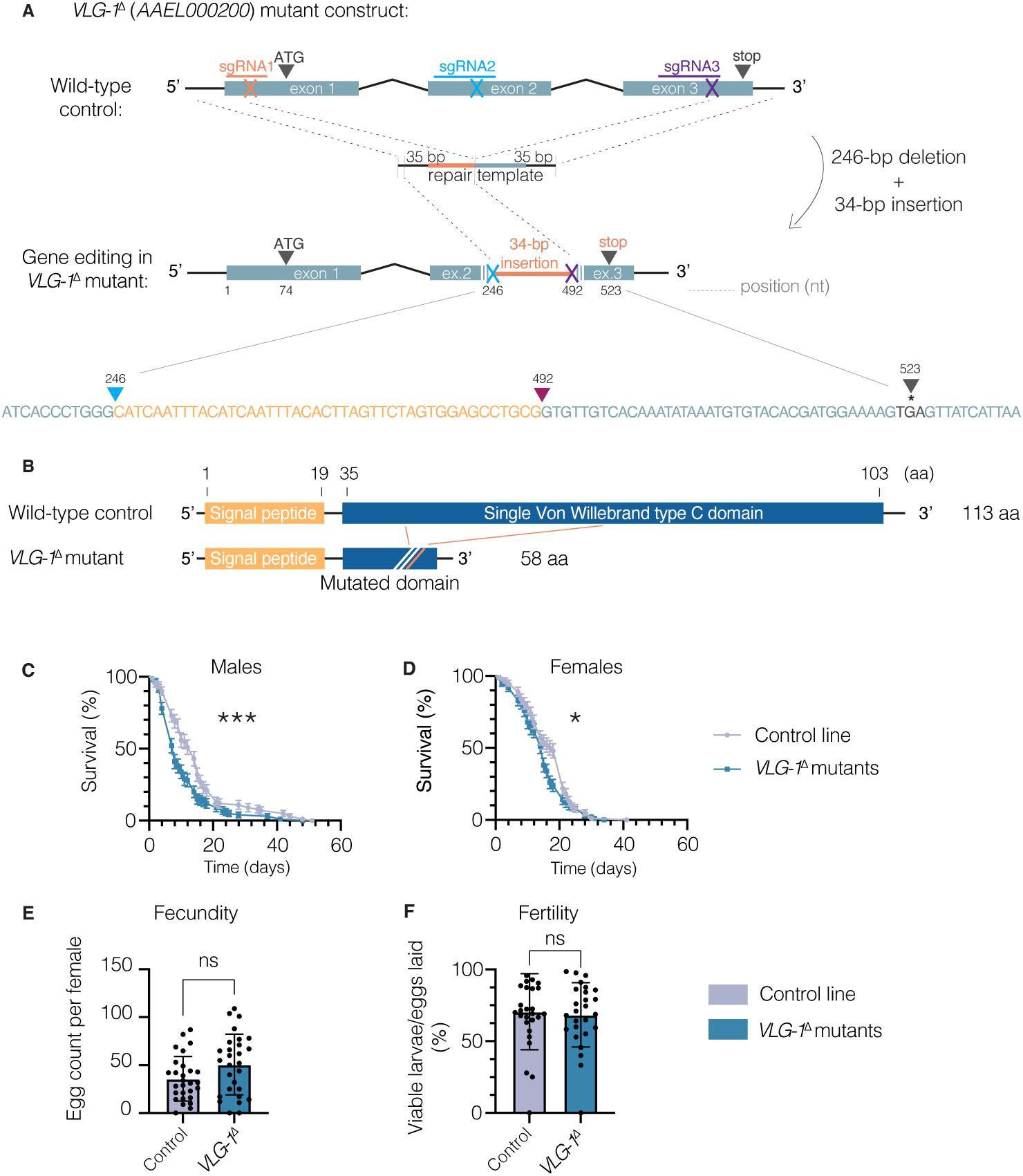
*VLG-1*^Δ^ mutants are viable and fertile without major fitness defects. (A) Structure of the *VLG-1* locus in the wild-type control and the *VLG-1*^Δ^ mutant lines. Exons are displayed as light blue boxes, connected by black segments representing introns. The positions and cut-sites of single-guide RNAs are depicted on each exon. (B) Structure of the VLG-1 protein in the wild-type control line and the *VLG-1*^Δ^ mutant line. (C-D) Survival curves of adult males (C) and females (D) from the wild-type control (grey) and *VLG-1*^Δ^ mutant (blue) lines in standard insectary conditions. Data represent mean and standard deviation of 4 replicates performed with 25 mosquitoes for each condition. *p<0.05; ***p<0.001 (Gehan-Breslow-Wilcoxon test). (E) Fecundity (number of eggs laid per individual blood-fed female for 7 days after a bloodmeal) in the *VLG-1*^Δ^ mutant and control lines. Data represents mean and standard deviation of 28 mosquitoes. *p<0.05; **p<0.01; ***p<0.001 (Mann-Whitney’s test). (F) Fertility (number of viable hatched larvae over the total number of eggs laid) in the *VLG-1*^Δ^ mutant and control lines. Data represent mean and standard errors of 26 mosquitoes. *p<0.05; **p<0.01; ***p<0.001 (chi-squared test).

To assess the impact of *VLG-1* absence on mosquito fitness, we monitored adult survival rates in standard insectary conditions. Mortality rates were slightly higher in the *VLG-1*^Δ^ mutant line compared to controls, particularly for males (Figure 3C-D). We also measured fecundity (*i.e.*, the number of eggs laid per blood-fed female; Figure 3E) and fertility (*i.e*., the number of viable larvae hatched over total number of eggs laid; Figure 3F) and found no differences between the *VLG-1*^Δ^ mutant and control lines. In sum, *VLG-1*^Δ^ mutants are viable and display no major fitness defects.

### The transcriptional landscape of *VLG-1*^Δ^ mutants is broadly altered

To investigate the overall impact of *VLG-1* loss and its potential link with virus infection in *Ae. aegypti*, we analyzed the midgut and body (carcass + head) transcriptomes of *VLG-1*^Δ^ mutant and control lines on days 2, 5, and 9 after a DENV-1 or mock bloodmeal. We detected transcripts from a total of ∼15,000 unique genes in midguts and ∼16,800 unique genes in bodies, representing 75% to 85% of all annotated genes depending on the samples and conditions.

In midguts, several hundreds of genes (ranging from 236 to 681) were significantly differentially expressed, defined by a fold change ≥ 2 and a p-value ≤ 0.05 between *VLG-1*^Δ^ mutants and controls (Figure 4A-B and Supplementary Figure S5). The highest number of differentially expressed genes (DEGs) was observed on day 2 after DENV exposure, with 380 up-regulated and 301 down-regulated genes. Overall, up to 4.5% of all detected genes were differently expressed between *VLG-1*^Δ^ mutants and controls in midguts. In the bodies, fewer DEGs were detected, but the highest number of DEGs was still detected 2 days after DENV exposure. These results suggest that *VLG-1* has a wide impact on biological processes, most prominently 2 days after a bloodmeal and especially in the presence of DENV.

**Figure 4.**
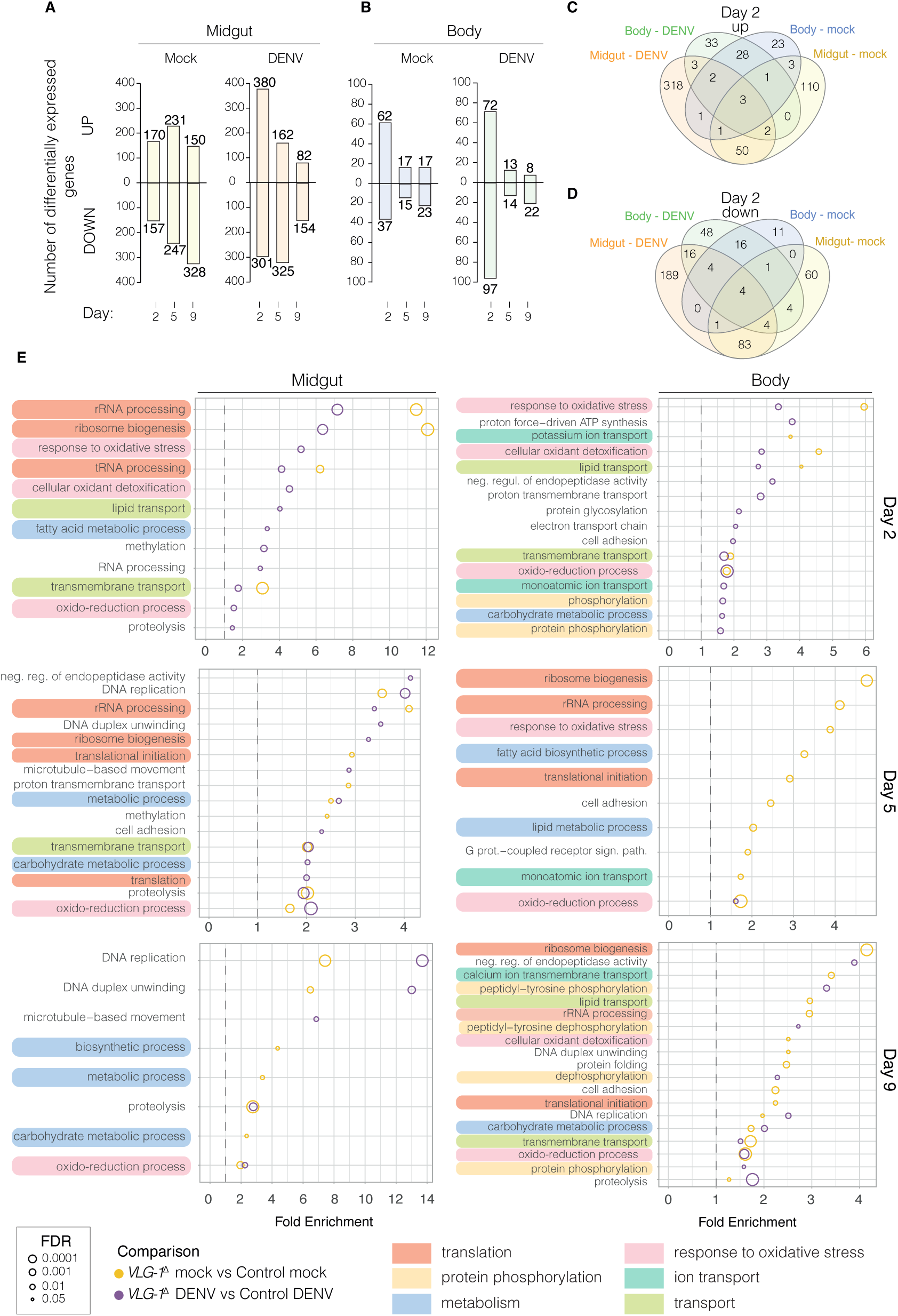
The transcriptome of *VLG-1*^Δ^ mutants is broadly altered. Female mosquitoes of the control and *VLG-1*^Δ^ mutant lines were offered a mock or infectious bloodmeal containing 5×10^6^ FFU/mL of DENV-1. On days 2, 5, and 9 post bloodmeal, 3 pools of 10 tissues (midguts or bodies) were collected and analyzed by RNA-seq. (A-B) Number of differentially expressed genes (DEGs) in *VLG-1*^Δ^ mutants compared to wild-type controls in midguts (A) and bodies (B) for both directions of change (up- or down-regulated). A gene was considered DEG when the absolute fold change was ≥ 2 and the adjusted p-value was ≤ 0.05. (C-D) Venn diagrams showing the overlap of up-regulated (C) and down-regulated (D) genes in *VLG-1*^Δ^ mutants compared to controls between all combinations of midguts, bodies, and bloodmeal type, on day 2 post bloodmeal. (E) Gene ontology (GO) enrichment of DEGs in *VLG-1*^Δ^ mutants compared to controls for both bloodmeal types, in midguts and bodies. Fold enrichment of each GO term is represented by a circle whose size is inversely proportional to the false discovery rate (FDR) of the enrichment score. GO terms with a similar biological function are identified with a color code and assigned a higher-order functional annotation. Correspondence between GO term names and IDs are listed in Supplementary Table 4.

*VLG-1*-dependent changes in gene expression occurred in the midgut and the rest of the body, but the overlap between DEGs in midguts and bodies was minimal (Figure 4C-D and Supplementary Figure S4). Only 18 and 34 up- or down-regulated transcripts (out of 684 and 592) were shared between both compartments, suggesting tissue-specific functions for *VLG-1*. Conversely, a noteworthy overlap of DEGs was detected between the mock and DENV bloodmeal conditions in both compartments, suggesting that *VLG-1*-dependent gene expression is only partially affected by virus infection.

To explore the biological functions of DEGs in *VLG-1*^Δ^ mutants, we examined their gene ontology (GO) annotations at the level of biological processes. We found that enriched GO terms in both midguts and bodies included mainly response to oxidative stress, translation regulation, and molecule transport (Figure 4E). Midgut-specific DEGs were mostly associated with RNA processing and broad metabolic processes, whereas most body-specific DEGs belonged to protein phosphorylation, protein modification, and ion transport categories. We did not specifically identify immune genes or pathways that were differentially expressed in *VLG-1*^Δ^ mutants. None of the genes previously reported to be involved in the activation or function of *CxVLG-1* (*Rel2*, *TRAF*, *Dicer2*, and *vir-1*) [26, 27] were DEGs in our dataset. This observation suggests that *Ae. aegypti VLG-1* and its *Culex* ortholog participate in different signaling pathways despite their close phylogenetic relatedness. However, these differences could also be explained by differences in experimental models. Previous studies on *CxVLG-1* primarily relied on *in vitro* approaches, which do not account for factors such as cell and tissue diversity or physiological processes like viral dissemination that occur during *in vivo* infections. Nevertheless, several DEGs were related to protein phosphorylation, particularly in DENV-exposed mosquitoes. These genes include several activators of immune pathways, such as *Pelle* and *Tube* in the Toll pathway, *Hop* in the JAK-STAT pathway, and *Tak* in the IMD pathway. Similarly, some DEGs identified in infected midguts and related to proteolysis were often associated with the Toll or IMD pathways, such as *CLIP* or *DREDD* genes. Thus, we cannot exclude a link between *Ae. aegypti VLG-1* and the canonical inducible immune pathways, although it would be distinct from previous observations in *Culex* studies. We also found an enrichment of DEGs involved in redox and oxidative stress response across tissues, timepoints and bloodmeal types. The anti-oxidative response was predominantly reduced in the bodies of the *VLG-1*^Δ^ mutants relative to the controls, suggesting that *VLG-1* limits cellular oxidation, possibly impacting antiviral host defense. Finally, we observed many enriched GO terms related to translation regulation, which might have a broad physiological impact.

### *VLG-1* slightly promotes systemic dissemination of DENV and ZIKV in *Ae. aegypti*

To investigate how the broad impact of *VLG-1* on the transcriptome functionally affects virus infection in *Ae. aegypti* mosquitoes, we performed experimental DENV-1 or ZIKV infections and analyzed infection prevalence (proportion of virus-positive tissues) and viral load (abundance of viral RNA) by RT-qPCR in individual tissues (midguts, carcasses, and heads). We selected timepoints representing key steps in the infection cycle: early midgut infection (day 2), systemic viral dissemination from the midgut to secondary organs (day 5), and head infection (day 9) (Figure 5 and Figure 6). Midgut infection prevalence is defined as the proportion of virus-positive midguts over the total number of blood-fed mosquitoes. Carcass infection prevalence is the proportion of virus-positive carcasses over the number of virus-positive midguts. Head infection prevalence is the number of virus-positive heads over the number of virus-positive carcasses. On days 7, 10, and 14 post bloodmeal, we measured viral titers in saliva samples collected from individual mosquitoes to assess virus transmission levels. Transmission efficiency was calculated as the proportion of virus-exposed mosquitoes with virus-positive saliva.

**Figure 5.**
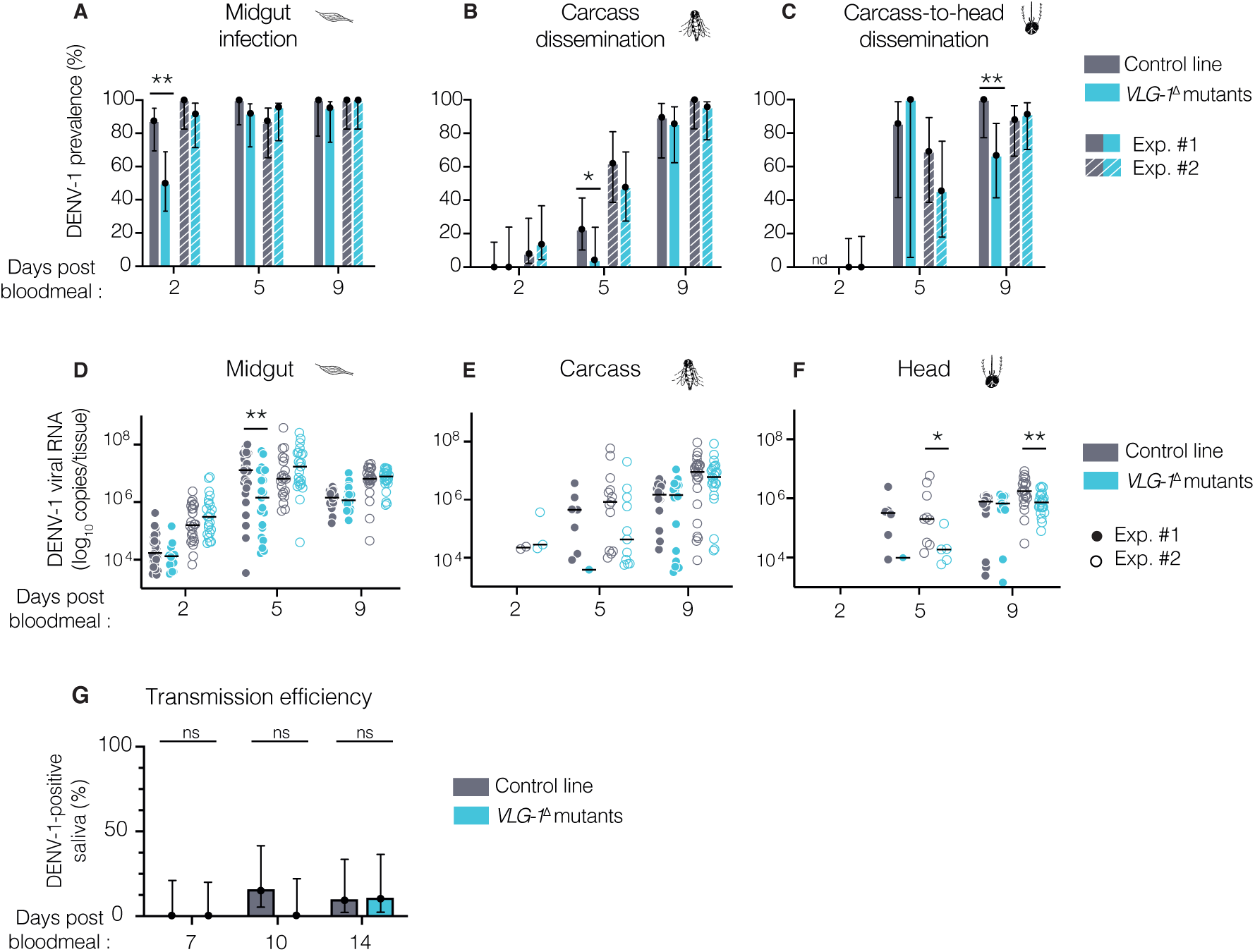
*VLG-1* slightly promotes systemic DENV dissemination in *Ae. aegypti*. (A-F) Female mosquitoes from the control (grey) and *VLG-1*^Δ^ mutant (blue) lines were offered an infectious bloodmeal containing 5×10^6^ FFU/mL of DENV-1. DENV-1 infection prevalence (A-C) and non-zero viral loads (D-F) were measured by RT-qPCR in the midgut, carcass, and head of individual mosquitoes on days 2, 5, and 9 post bloodmeal. (A-C) Midgut infection prevalence was calculated as the number of virus-positive midguts over the total number of virus-exposed mosquitoes. Carcass dissemination prevalence was calculated as the number of virus-positive carcasses over the number of virus-positive midguts. Carcass-to-head dissemination prevalence was calculated as the number of virus-positive heads over the number of virus-positive carcasses. (D-F) Each dot represents an individual tissue. The horizontal black lines represent the median values. *p<0.05; **p<0.01; ***p<0.001 (Mann-Whitney’s test). (G) Saliva samples from virus-exposed mosquitoes were collected on days 7, 10, and 14 after exposure and infectious virus particles in the saliva were detected by focus-forming assay. In (A-C) and (G), vertical error bars represent the 95% confidence intervals of the proportions. *p<0.05; **p<0.01; ***p<0.001 (chi-squared test). In (A-F), data from two experimental replicates, analyzed and displayed separately because a significant experiment effect was detected, are plotted using two shades of the same color.

**Figure 6.**
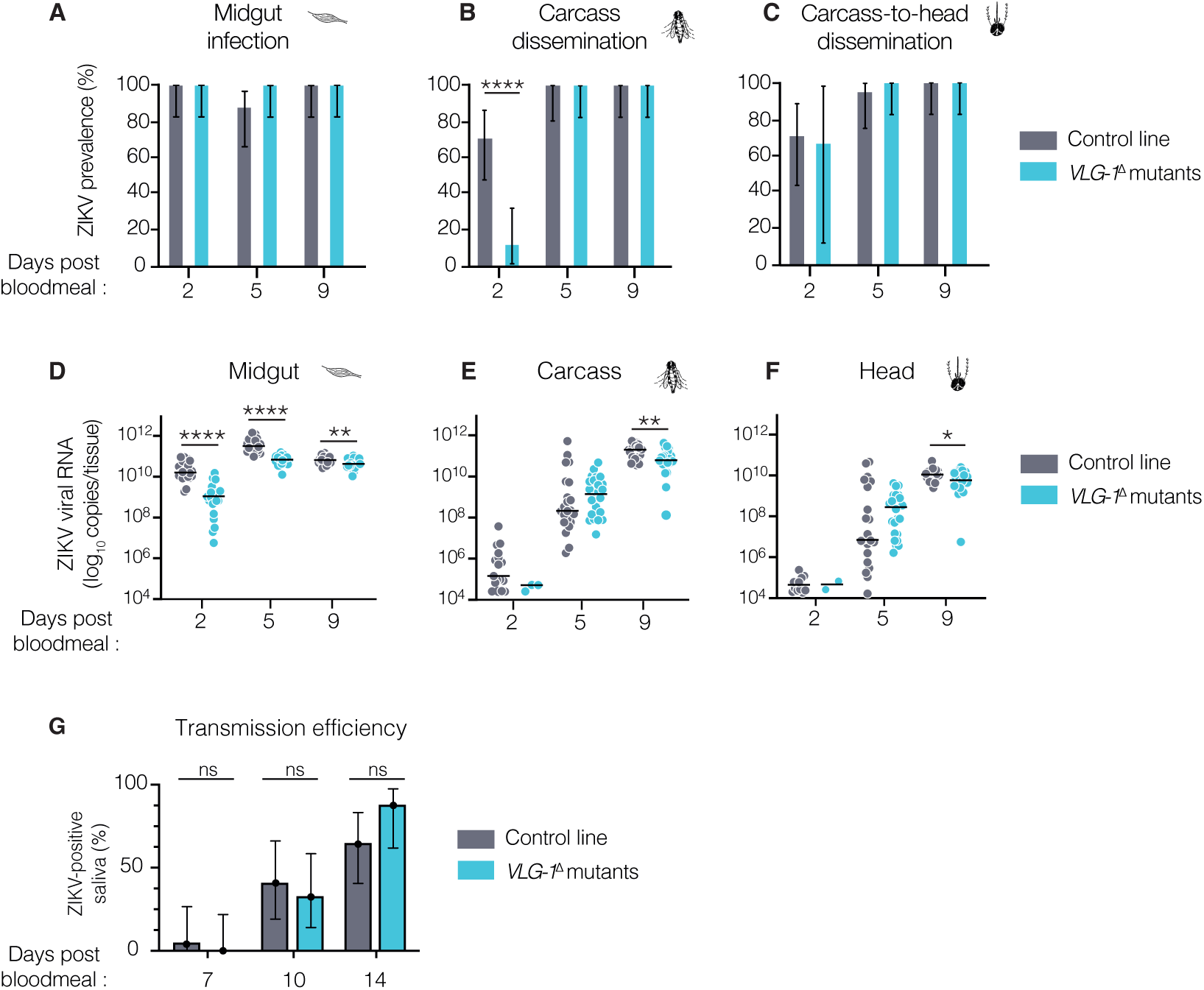
*VLG-1* slightly promotes systemic ZIKV dissemination in *Ae. aegypti*. (A-F) Female mosquitoes from the control (grey) and *VLG-1*^Δ^ mutant (blue) lines were offered an infectious bloodmeal containing 5×10^5^ PFU/mL of ZIKV. ZIKV infection prevalence (A-C) and non-zero viral loads (D-F) were measured by RT-qPCR in the midgut, carcass, and head of individual mosquitoes on days 2, 5, and 9 post bloodmeal. (A-C) Midgut infection prevalence was calculated as the number of virus-positive midguts over the total number of virus-exposed mosquitoes. Carcass dissemination prevalence was calculated as the number of virus-positive carcasses over the number of virus-positive midguts. Carcass-to-head dissemination prevalence was calculated as the number of virus-positive heads over the number of virus-positive carcasses. (D-F) Each dot represents an individual tissue. The horizontal black lines represent the median values. *p<0.05; **p<0.01; ***p<0.001 (Mann-Whitney’s test). (G) Saliva samples from virus-exposed mosquitoes were collected on days 7, 10, and 14 after exposure and infectious virus particles in the saliva were detected by focus-forming assay. In (A-C) and (G), vertical error bars represent the 95% confidence intervals of the proportions. *p<0.05; **p<0.01; ***p<0.001 (chi-squared test).

Upon DENV-1 infection, we found that the dynamics of systemic dissemination slightly differed between *VLG-1*^Δ^ mutants and wild-type controls (Figure 5A-F). All the statistically significant differences that we detected indicated that virus dissemination was slower in *VLG-1*^Δ^ mutants. This effect was consistent but manifested differently in two experimental replicates. In the first experimental replicate, we found that infection prevalence in the midgut (day 2), carcass (day 5) and head (day 9) was lower in *VLG-1*^Δ^ mutant mosquitoes (Figure 5A-C). In the second experimental replicate, we detected decreased viral loads in the *VLG-1*^Δ^ mutant midguts on day 5, and in heads on days 5 and 9 (Figure 5D-F). Such variation between experimental replicates presumably reflects minor uncontrolled variation in the bloodmeal titers that result in slightly different infection dynamics. Finally, we found no difference in DENV transmission efficiency between wild-type and *VLG-1*^Δ^ mutant mosquitoes (Figure 5G).

Next, we performed a similar set of experiments with ZIKV and confirmed *VLG-1*’s proviral effect on virus dissemination (Figure 6). Two days after the infectious bloodmeal, we found a significant decrease in infection prevalence in the carcass, where only 12% of the *VLG-1*^Δ^ mosquitoes harbored ZIKV RNA compared to 70% of the control mosquitoes (Figure 6A-C). In midguts, we consistently found a decrease in viral loads (∼10-fold) in *VLG-1*^Δ^ mutants at all three timepoints (Figure 6D). In carcasses and heads, viral loads were 5- to 10-fold lower in *VLG-1*^Δ^ mutants 9 days post bloodmeal (Figure 6E-F). Similar to DENV, we found no detectable difference in virus transmission efficiency between wild-type and *VLG-1*^Δ^ mutant mosquitoes (Figure 6G).

Our data demonstrate that *VLG-1* slightly promotes flavivirus dissemination across the mosquito’s body but does not seem to significantly impact virus transmission. Of note, we estimated virus transmission efficiency with standard salivation assays that potentially underestimate vector competence compared to live-host transmission assays [38], which might have limited our ability to detect differences in transmission efficiency between the *VLG-1*^Δ^ mutant and control lines. Together, these results show that *VLG-1* lacks any antiviral activity and rather exerts a modest proviral effect during flavivirus infection of *Ae. aegypti in vivo*.

### *VLG-1* and *VLG-2* have non-additive proviral effects on DENV in *Ae. aegypti*

The finding of *VLG-1’*s proviral effect prompted us to test whether its paralog *VLG-2* could share similar properties in *Ae. aegypti*. Using RNAi-mediated knockdown, we depleted *VLG-2* transcripts in adult *VLG-1*^Δ^ mutants or control mosquitoes. Two days after injection of double-stranded RNA (dsRNA) targeting *VLG-2* or *Luciferase* (as a control), mosquitoes were exposed to a DENV-1 infectious bloodmeal and their heads collected 7 days later. Consistent to previous results, we found that infection prevalence in heads was lower for *VLG-1*^Δ^ mutants than for wild-type mosquitoes upon control dsRNA injection (Supplementary Figure S3A). Head infection prevalence was also lower in wild-type mosquitoes depleted in *VLG-2* transcripts, revealing a proviral role for *VLG-2*. Finally, head infection prevalence was not further reduced in mosquitoes that were depleted for both *VLG-1* and *VLG-2* transcripts (Supplementary Figure S3A). Additionally, we did not detect differences in viral loads between any of the experimental treatments (Supplementary Figure S2B). On the day of the infectious bloodmeal, we tested *VLG-2* gene knockdown efficiency and found a strong reduction in transcript abundance for both isoforms (*VLG-2-RA* and -*RB*) in all conditions (Supplementary Figure S3C-D). Together, these results indicate that *VLG-1* and *VLG-2* exert non-additive proviral effects on DENV infection in *Ae. aegypti*.

## DISCUSSION

In this study, we identified the *Ae. aegypti* gene *AAEL000200* as a Culicinae-specific *Vago*-like gene that we renamed *AaeVLG-1*. We generated a *VLG-1* mutant line of *Ae. aegypti* that displayed a slight reduction in lifespan but remained fully viable and fertile. Our tissue-specific transcriptomic analysis showed a broad remodeling of gene expression in *VLG-1*^Δ^ mutants. Additionally, we found that during DENV and ZIKV infection of *Ae. aegypti in vivo*, *VLG-1* exerted a subtle proviral role by enhancing virus dissemination, but not virus transmission. Our *in vivo* approach offers the first dynamic insight into *VLG-1* function during flavivirus infection in *Ae. aegypti*, uncovering compartment-specific and time-dependent effects of this gene. Together, this work challenges the assumed universal nature of the antiviral function of *Vago*-like genes in arthropods.

Our transcriptomic analysis revealed that the loss of *VLG-1* interferes with a wide range of biological pathways. Notably, canonical immune pathways were not significantly impacted by *VLG-1* loss of function. Amongst the most significantly altered processes in *VLG-1*^Δ^ mutants was the response to oxidative stress. Pro-oxidative processes were up-regulated and anti-oxidative processes were down-regulated in the *VLG-1*^Δ^ mutants, suggesting that *VLG-1* confers protection against oxidative stress. Hijacking of oxidative stress by viruses has been reported to facilitate their genome replication [39-42]. Additionally, oxidative stress can also contribute to the cellular antiviral response [43-45]. Thus, modulation of the oxidative stress response by *VLG-1* could contribute to its proviral effect and explain the shorter lifespan of *VLG-1*^Δ^ mutants.

The induction mechanism of *VLG-1* remains to be elucidated. In *Culex* mosquitoes, *CxVLG-1* induction depends on a NF-κB Rel-binding site [27]. We ran a promoter analysis to identify transcription factor (TF) binding motifs in the promoter sequence of *VLG-1* (Supplementary Figure S6 and Supplementary Table 6). Importantly, we did not identify classical immune TF binding motifs, such as NF-κB motifs. In contrast, we identified TF binding motifs specific to signaling pathways involved in cell cycle regulation, apoptosis, and redox stress response. This observation is consistent with the hypothesis that *VLG*-1’s modest proviral activity in *Ae. aegypti* is not associated with canonical immune pathways but rather with stress response processes.

Mechanistic insights into *VLG-1*’s mode of action in *Ae. aegypti* remain to be investigated. CxVLG-1 is secreted extracellularly in *Culex*-derived cells [26], and Vago-like proteins are presumed to be secreted in several other insect species [25, 32]. The AaeVLG-1 protein sequence contains a secretion signal peptide, but experimental evidence of extracellular localization is lacking. Technical limitations such as minute protein amounts in the mosquito hemolymph, low sensitivity of detection, and lack of adequate controls prevented us from assessing the extracellular presence of VLG-1 *in vivo* by immunoblotting. Mass spectrometry analysis of the hemolymph protein content may be required to confirm VLG-1 secretion in the extracellular environment.

Our evolutionary analyses of *Vago*-like gene homologs in dipteran insects showed that both *VLG* paralogs have been retained and maintained under slightly relaxed selective pressure since the Culicinae diversification 150 million years ago. This indicates that they did not undergo pseudogenization (*i.e*., accumulation of deleterious mutations resulting in a non-functional gene sharing high sequence identity with the ancestral form). Our knockdown experiments also revealed a proviral effect of *AaeVLG-2*, but this remains to be more comprehensively investigated. The results of our evolutionary analysis do not support the hypothesis of neofunctionalization of *VLG-1* following its duplication from *VLG-2*. We found that purifying selection remained the predominant mode of evolution for both paralogs after the duplication in the Culicinae. Neofunctionalization is typically associated with relaxed purifying selection, including sites evolving under positive selection and diversification [46], as well as asymmetry in ω following the duplication event [47]. Our results are more consistent with subfunctionalization, whereby each paralog retains a subset of its original ancestral function. Under a subfunctionalization scenario, higher ω is expected in the daughter lineages compared to the parental lineage [47]. Moreover, *VLG-2* knockdown in *VLG-1*^Δ^ mutants resulted in a similar phenotype to *VLG-2* knockdown in wild-type controls and control knockdown in *VLG-1*^Δ^ mutants, suggesting a functional co-dependency of *VLG-2* and *VLG-1*, where both paralogs would provide their proviral activity jointly in *Ae. aegypti*. Subfunctionalization can also occur via specialization, a process in which paralogs divide into various areas of specialty, such as tissue-specificity, rather than function [48]. Additional evidence is needed to support a subfunctionalization scenario for *Vago*-like genes in *Ae. aegypti*.

In conclusion, our study provides a dynamic view of *VLG-1* function during flavivirus dissemination in *Ae. aegypti*. Unexpectedly, this *in vivo* work reveals a subtle proviral activity of *VLG-1* that is both time-sensitive and tissue-specific, an aspect previously overlooked in *in vitro* studies. Although the modest proviral effect of *VLG-1* does not seem to significantly influence vector competence, our findings challenge the notion that genes of the *Vago* family are conserved antiviral factors in arthropods and question their designation as antiviral cytokines. We anticipate that our newly generated *VLG-1*^Δ^ mosquito mutant line will serve as a valuable tool to investigate the function of *VLG-1* in *Ae. aegypti*. This work underscores the importance of *in vivo* research for identifying and characterizing the biological roles of pro- and antiviral factors that govern the ability of *Ae. aegypti* mosquitoes to transmit arboviruses. This fundamental understanding of mosquito-arbovirus interactions will be critical to the development of new strategies aiming to reduce the burden of arboviral diseases [49].

## METHODS

### Virus strains

DENV-1 strain KDH0026A was originally isolated in 2010 from the serum of a patient in Kamphaeng Phet, Thailand [50]. ZIKV strain Kedougou2011 was originally isolated in 2011 from a mosquito pool in Kedougou, Senegal [51]. Viral stocks were prepared in C6/36 *Aedes albopictus* cells as previously described [52].

### Mosquitoes

Experiments were conducted with a previously described isofemale line of *Ae. aegypti* called Jane [19, 53]. Mosquitoes were reared in controlled conditions (28°C, 12-hour light/12-hour dark cycle and 70% relative humidity). For experiments, eggs were hatched synchronously in a SpeedVac vacuum device (Thermo Fisher Scientific) for 45 minutes. Larvae were reared in plastic trays containing 1.5 L of dechlorinated tap water and fed a standard diet of Tetramin (Tetra) fish food at a density of 200 larvae per tray. After emergence, adults were kept in BugDorm-1 insect cages (BugDorm) with permanent access to 10% sucrose solution.

### CRISPR/Cas9-mediated gene editing

#### sgRNA design and synthesis

A *VLG-1* mutant line and wild-type “sister” line were derived from the 26^th^ generation of the Jane isofemale line. Gene editing was performed using CRISPR/Cas9 technology as previously described [54]. The single-guide RNAs (sgRNAs) were designed using CRISPOR [55] by searching for 20-bp sgRNAs with the NGG protospacer-adjacent-motif (PAM). To reduce chances of off-target mutations, only sgRNAs with off-target sites with at least four mismatches were selected. Three sgRNAs were selected with cut sites respectively located upstream of the start codon, in the middle of the *VLG-1* gene within the second exon, and upstream of the stop codon. Since the *VLG-1* locus is only 471-bp (including introns), a single-stranded oligodeoxynucleotide (ssODN) repair template was provided to delete the entire gene. The ssODN repair template included two 35-bp homology arms matching the sequence upstream from the cut site of the first sgRNA (x1_30rev) and downstream from the cut site of the third sgRNA (x3_67rev) to facilitate excision of the *VLG-1* gene. The ssODN repair template was synthesized and PAGE-purified commercially (Sigma-Aldrich). Single-guide RNAs were synthetized with the with MEGAscript T7 *in vitro* transcription kit (Ambion) and purified with the MEGAclear kit (Invitrogen).

#### Embryonic microinjections

*Ae. aegypti* embryos were injected with a microinjection mix containing 402.5 ng/µL SpCas9 protein (New England Biolabs), 40 ng/µL of each of three sgRNAs (x1_30rev, x2_6rev, x3_67rev), and 125 ng/µL of the ssODN repair template suspended in molecular grade water. The microinjection of *Ae. aegypti* embryos was performed using standard protocols [56]. *Ae. aegypti* adult females were bloodfed with commercial rabbit blood (BCL) via an artificial membrane feeding system (Hemotek). Three days post bloodmeal, females were transferred to egg-laying vials and oviposition was induced by placing mosquitoes into dark conditions for 15 min. Embryos were injected 30-60 min post oviposition. Embryos were hatched in water 3 days post injection and individual pupae placed into vials containing a small amount of water to isolate and screen adults for mutations before mating could occur.

#### Mutation isolation and line creation

Individual virgin adult G_0_ mosquitoes were screened for mutations by PCR to amplify the *VLG-1* gene from DNA extracted from a single leg (see Genotyping below). The amplified region was 793 bp and deletions were screened for on a 2% agarose gel. If large deletions were detected, the corresponding mosquito was mated with wild-type mosquitoes of the opposite sex and progeny screened for inheritance of the mutation. Sanger sequencing was then performed to characterize the edit. A large deletion of ∼200 bp was identified in a G_0_ female that was subsequently placed in a cage with 3 wild-type males for mating, blood feeding, and egg laying. The G_1_ eggs were hatched in water 5 days post laying and individual pupae isolated into vials containing a small amount of water to isolate and screen adults for mutations before mating could occur. G_1_ progeny was screened for the deletion by PCR to confirm heritability of the mutation. Four G_1_ males (heterozygous for the mutation) were then crossed with 50 wild-type females. Next, 11 G_2_ male heterozygotes were crossed with 23 wild-type females. Finally, 14 G_3_ males and 33 G_3_ females heterozygous at the mutation site were crossed with each other. G_4_ adults were sorted into homozygous mutants (establishing the *VLG-1*^Δ^ mutant line) and homozygous wild types (establishing the control “sister” line). The *VLG-1*^Δ^ mutant line was established with 9 G_4_ males and 24 G_4_ females, while the control line was established with19 males and 29 females. Sequencing of *VLG-1*^Δ^ individuals using Sequencing primer F (Supplementary Table 5) revealed that the deletion spanned 246 bp of the wild-type *VLG-1* sequence starting at the cut site of the sgRNA in the middle of the gene (x2_6rev), 170 bp downstream of the still intact start codon, and ending at the cut site of the third sgRNA (x3_67rev), 49 bp upstream of the stop codon. However, the mutation also contained a 34-bp insertion of the upstream 35-bp homology arm of the repair template in-between the sgRNA cut sites, resulting in a PCR product 212-bp shorter than the wild-type PCR product, matching what was visualized on gels during screening.

#### Genotyping

Genomic DNA was extracted from single legs of individual mosquitoes using DNAzol DIRECT (DN131, Molecular Research Center, Inc.). To obtain legs from live mosquitoes, pupae were placed in vials containing a small volume of water and sealed with a cotton plug (Flugs, Genesee). After adult emergence, the water was drained and vials placed on ice for anesthesia. Single legs were collected using forceps and placed in a 2-mL screw-top plastic tube containing ∼20 1-mm glass beads (BioSpec) and 200 µL of DNAzol DIRECT. Mosquitoes were then placed back into the vials to remain isolated and unmated until genotyping via PCR. The legs were homogenized for 30 sec at 6,000 revolutions per minute (rpm) in a Precellys 24 tissue homogenizer (Bertin Technologies), briefly centrifuged, and then placed at room temperature (20-25°C) for immediate use. PCR was performed using DreamTaq DNA Polymerase (EP0701, Thermo-Fisher Scientific) based on manufacturer’s instructions, using Genotyping primers (Supplementary Table 5). Approximately 0.6 µL of the DNAzol DNA extract from leg tissue was used in 19 µL of DreamTaq PCR master mix. The PCR conditions were as follows: initial denaturation at 95°C for 3 min, 40 cycles of amplification (denaturation at 95°C for 30 sec, annealing at 59°C for 15 sec, and extension at 72°C for 30 sec), and a final extension step at 72°C for 5 min. Amplicons were purified (MinElute PCR purification kit, Qiagen) and subsequently sequenced (Eurofins).

### Evolutionary analyses

#### Gene phylogeny

Using the protein sequence of *AaeVLG-1* (*AAEL000200;* RefSeq accession number XP_001658930.1) and *AaeVLG-2* (*AAEL000165;* RefSeq accession number XP_001658929.1) as queries, we performed a BLASTP against the NCBI non-redundant protein database to extract homologous genes present in the *Drosophila* genus and Culicidae family. Only genes present in the reference sequence (RefSeq) were considered in the final dataset of 62 homologous genes (Supplementary Table 2). Then, input coding sequences were aligned with respect to their codon structure using MACSE v2.06 [57] and the protein alignment was used as input for IQ-tree2 [58] to infer the phylogenetic relationships of the *Vago*-like gene homologs. The substitution model WAG+I+G4 was the best fit model based on the Bayesian Information Criterion (BIC) and the maximum-likelihood phylogenetic tree was generated with 1,000 ultra-fast bootstrap iterations. The phylogenetic tree of *Vago*-like genes was rooted using *Drosophila* sequences and visualized using iTOL [59].

#### Evolutionary rate

To investigate the evolutionary rates of *Vago*-like gene coding sequences, the CODEML tool from the PAML package [60] was used to detect variations of the ratio of non-synonymous to synonymous substitutions (ω) as a proxy for the variation in selective pressure, following the guide for user good practices [61]. CODEML was configured to use the branch model, which assumes different ω parameters for different branches in the phylogeny [62, 63]. Three tests were conducted by designating different branches as the foreground: (i) *VLG-1* branch, (ii) both *VLG* and *VLG-1* branches, (iii) *VLG-1*, Anophelinae *VLG* and Culicinae *VLG* branches. Comparison of the branch model to the null model was performed through a likelihood-ratio test (Supplementary Table 3).

### Mosquito fitness assays

#### Survival

Five to seven days after adult emergence, males and females were sorted and transferred in 1-pint carton boxes with permanent access to 10% sucrose solution at 28°C and 70% relative humidity. Mortality was scored daily. Four replicate boxes containing 25 mosquitoes each were used for each experiment.

#### Fecundity

Five- to seven-day-old females were blood fed and transferred to individual vials containing a humid blotting paper for egg laying with access to 10% sucrose solution. After 7 days, eggs deposited on the blotting paper were counted under a binocular magnifier. Fecundity was defined as the number of eggs laid per blood-fed female.

#### Fertility

The aforementioned blotting papers air dried for a week. Eggs were then hatched synchronously in a SpeedVac vacuum device (Thermo Fischer Scientific) for 45 min. Larvae were transferred to individual vials containing tap water and with Tetramin fish food, and viable larvae were enumerated three days later. Fertility was defined as the number of viable larvae over the total number of laid eggs per blood-fed female.

### Mosquito infectious bloodmeals

Experimental infections of mosquitoes were performed in a biosafety level-3 containment facility, as previously described [52]. Shortly, 5- to 7-day-old female mosquitoes were deprived of 10% sucrose solution 20 hours prior to being exposed to an artificial infectious bloodmeal containing 5×10^6^ FFU/mL of DENV-1 or 5×10^5^ PFU/mL of ZIKV. The infectious bloodmeal consisted of a 2:1 mix of washed rabbit erythrocytes (BCL) supplemented with 10 mM adenosine triphosphate (Sigma) and viral suspension supplemented with Leibovitz’s L-15 medium (Gibco; described below). Mosquitoes were exposed to the infectious bloodmeal for 15 min through a desalted pig-intestine membrane using an artificial feeder (Hemotek Ltd) set at 37°C. Fully blood-fed females were sorted on ice and incubated at 28°C, 70% relative humidity and under a 12-hour light-dark cycle with permanent access to 10% sucrose solution.

### Gene expression and viral load quantification

Mosquito tissues were dissected in 1× phosphate-buffered saline (PBS), and immediately transferred to a tube containing 400 µL of RA1 lysis buffer from the Nucleospin 96 RNA core kit (Macherey-Nagel) and ∼20 1-mm glass beads (BioSpec). Samples were homogenized for 30 sec at 6,000 rpm in a Precellys 24 grinder (Bertin Technologies). RNA was extracted and treated with DNase I following the manufacturer’s instructions. Viral RNA was reverse transcribed and quantified using a TaqMan-based qPCR assay, using virus-specific primers and 6-FAM/BHQ-1 double-labelled probe (Supplementary Table 5). Reactions were performed with the GoTaq Probe 1-Step RT-qPCR System (Promega) following the manufacturer’s instructions. Viral RNA levels were determined by absolute quantification using a standard curve. The limit of detection was of 40 copies of viral RNA per microliter. Transcript RNA levels were normalized to the housekeeping gene encoding ribosomal protein S 17 (*RPS17*), and expressed as 2^-dCt^, where dCt = Ct*_Gene_* – Ct*_RPS17_*.

### Virus titration

#### Focus-forming assay (FFA)

DENV infectious titers were measured by standard FFA in C6/36 cells. Cells were seeded at a density of 5×10^4^ cells/well in a 96-well plate 24 hours before inoculation. Serial sample dilutions were prepared in Leibovitz’s L-15 medium (Gibco) supplemented with 0.1% penicillin/streptomycin (pen/strep; Gibco ThermoFisher Scientific), 2% tryptose phosphate broth (TBP; Gibco Thermo Fischer Scientific), 1× non-essential amino acids (NEAA; Life Technologies) and 2% fetal bovine serum (FBS; Life Technologies). Cells were inoculated with 40 μL of sample. After 1 hour of incubation at 28°C, the inoculum was replaced with 150 μL of overlay medium (1:1 mix of Leibovitz’s L-15 medium supplemented with 0.1% pen/strep, 2% TPB, 1× NEAA, 2× Antibiotic-Antimycotic [Life Technologies], 10% FBS and 2% carboxyl methylcellulose) and incubated for 5 days at 28°C. Cells were fixed for 30 min in 3.6% paraformaldehyde (PFA; Sigma-Aldrich). Cells were then washed three times with PBS 1×, and permeabilized for 30 min with 50 μL of PBS 1×; 0.3% Triton X-100 (Sigma-Aldrich) at room temperature (20-25°C). The cells were washed three times in PBS 1× and incubated for 1 hour at 37°C with 40 μL of mouse anti-DENV complex monoclonal antibody MAB8705 (Merck Millipore) diluted 1:200 in PBS 1×; 1% bovine serum albumin (BSA) (Interchim). After another three washes in PBS, cells were incubated at 37°C for 30 min with 40 μL of an Alexa Fluor 488-conjugated goat anti-mouse antibody (Life Technologies) diluted 1:500 in PBS 1×; 1% BSA. After three washes in PBS 1× and a final wash in water, infectious foci were counted under a fluorescent microscope (Evos) and converted into focus-forming units/mL (FFU/mL).

#### Plaque assay

ZIKV infectious titers were measured by plaque assay in Vero E6 cells. Cells were seeded in 24-well plates at a density of 150,000 cells/well 24 hours before inoculation. Ten-fold sample dilutions were prepared in Dulbecco’s Modified Eagle Medium (DMEM) with 2% FBS, 1% pen/strep, 4× Antibiotic-Antimycotic and cells were incubated with 200 μL of inoculum. After 1 hour at 37°C, the inoculum was replaced with DMEM supplemented with 2% FBS, 1% pen/strep, 4× Antibiotic-Antimycotic and 0.8% agarose. Cells were fixed with 3.6% PFA after 6 days and plaques were counted manually after staining with 0.1% crystal violet (Sigma).

### Salivation assay

Mosquitoes were anesthetized with triethylamine (≥99%, Sigma-Aldrich) for 10 min and their legs were removed. The proboscis of each female was inserted into a 20-µL pipet tip containing 10 µL of FBS for 30 min at room temperature (20-25°C). Saliva-containing FBS was expelled into 90 µL of Leibovitz’s L-15 medium supplemented with 0.1% pen/strep, 2% TPB, 1× NEAA and 4× Antibiotic-Antimycotic. Virus presence in saliva samples was determined by virus titration after 5 days of amplification in C6/36 cells. Transmission potential was assessed qualitatively based on the presence or absence of infectious virus.

### Transcriptome analysis

#### Library preparation and mRNA sequencing

Total RNA extracts from pools of 10 tissues were isolated with TRIzol (Invitrogen) as previously described [64] and treated with DNA-free kit (Invitrogen, AM1906) following the manufacturer’s instructions. The quality of the samples was assessed using a BioAnalyzer RNA Nano kit (Agilent Technologies). RNA libraries were built using an Illumina Stranded mRNA library Preparation Kit (Illumina) following the manufacturer’s protocol depending on the insert size required. Of note, to obtain 300-bp inserts, all the samples were eluted for 2 minutes at 80°C after polyA capture, instead of the 8-min fragmentation at 94°C recommended by the supplier. Sequencing was performed on two lanes 10B300 of NovaSeqX (Illumina) by Novogene.

#### Bioinformatics

Raw RNA-seq reads were cleaned of adapter sequences and low-quality sequences using cutadapt version 2.10 [65] with options “-m 25 -q 30 -O 6 --trim-n --max-n 1”. Gene expression quantification was performed using salmon version 1.9.0 [66]. First, the *Ae. aegypti* reference transcriptome (downloaded from VectorBase (release 66) at https://vectorbase.org/common/downloads/release-66/AaegyptiLVP_AGWG/fasta/data/) was indexed along with its corresponding genome using the “--decoys” option. Transcript expression was then quantified for each sample using the “-l A” option and summarized at the gene level using the “--geneMap” parameter [67, 68]. Gene expression data was imported into R version 4.3.2 [69] using the tximport package [70]. The normalization and dispersion estimation were performed with DESeq2 [71] using the default parameters and statistical tests for differential expression were performed applying the independent filtering algorithm. For each tissue (bodies and midguts) at each time point (days 2, 5, and 9), a generalized linear model was set to test for the mutation effect on gene expression, separately for infected and non-infected mosquitoes. For each pairwise comparison, raw p-values were adjusted for multiple testing according to the Benjamini and Hochberg procedure [72] and genes with an adjusted p-value lower than 0.05 and an absolute fold-change higher than 2 were considered differentially expressed.

### Gene set enrichment analysis

Gene set enrichment analysis was performed using Fisher’s statistical test for the over-representation in differentially expressed genes. *Ae. aegypti* gene ontology (GO) annotations [73] were retrieved from the VectorBase website (version 66). Only gene sets with a false discovery rate (FDR) lower than 0.05 were considered significantly enriched in differentially expressed genes.

### Gene knockdown assay

Double-stranded RNA (dsRNA) targeting *AaeVLG-2* (AAEL000165) was *in vitro* transcribed from T7 promoter-flanked PCR products using the MEGAscript RNAi kit (Life Technologies). To obtain the PCR products, a first PCR was performed on genomic DNA extracted from wild-type mosquitoes using the previously described Pat-Roman DNA extraction protocol [74]. The T7 sequence was then introduced during a second PCR using T7 universal primers that hybridize to short GC-rich tags introduced to the PCR products in the first PCR (Supplementary Table 5). dsRNA targeting *Luciferase* (as a negative control) was synthesized using T7 promoter-flanked PCR products generated by amplifying a *Luciferase*-containing plasmid with T7-flanked PCR primers with the MEGAscript RNAi kit (Life Technologies) (Supplementary Table 5). dsRNA was resuspended in RNase-free water to reach a final concentration of 10 mg/mL. Five- to seven-day-old females were anesthetized on ice and injected intrathoracically with 1 µg (in a volume of 100 nL) dsRNA suspension using a Nanoject III apparatus (Drummond). After injection, mosquitoes were incubated for 2 days at 28°C before the infectious bloodmeal. The knockdown efficiency was estimated by RT-qPCR on the day of the bloodmeal as (1 – ddCt)*100, where ddCt = (mean (2^-dCt^ in ds*VLG-2* condition)) / (mean (2^-dCt^ in ds*Luciferase* condition)), and dCt = Ct*_VLG-2_* – Ct*_RPS17_*.

### Western blotting

Five female mosquitoes were collected in 250 μL of 2× RIPA buffer complemented with protease inhibitor (Complete 1×, Roche) in tubes containing ∼20 1-mm glass beads (BioSpec). Samples were homogenized for 30 sec at 6,000 rpm in a Precellys 24 grinder (Bertin Technologies). Lysates were clarified by centrifugation at 14,000 rpm for 5 min at 4°C and kept on ice. Fifty microliters of lysate were heated at 95°C with 50 μL of Laemmli buffer for 5 min. Twenty microliters of denatured samples were loaded on a PROTEAN TGX 4-20% stain-free precast gel (Biorad) in 1× Tris-Glycine-SDS running buffer (Alfa Aesar). Transfer on a nitrocellulose membrane was done using a Trans-Blot Turbo transfer pack (Biorad) for 30 min at 25 V. The membrane was then incubated in PBS 1×-Tween 0.1%-powdered milk (Régilait) 5% (PBST-milk) for 1 hour. Incubation with the primary antibody (rabbit anti-CxVLG-1 (GenScript) generated in [26], 1:2,000 in PBST-milk) was done for 1 hour at room temperature (20-25°C) before washing three times for 5 min in PBST. The anti-CxVLG-1 antibody targets the C-terminal sequence CEKIKQDLTKDYPE which is located within the deleted region in the *VLG-1*^Δ^ mutant sequence. The membrane was then incubated in the secondary antibody (donkey anti-rabbit, ab216779, 1:20,000 in PBST-milk) for 1 hour at room temperature. After three washes of 5 min in PBST, the membrane was imaged on an Odyssey LICOR imager.

### Promoter analysis

To analyze the presence of transcription factor (TF) binding motifs in the promoter of *VLG-1*, we used MoLoTool (https://molotool.autosome.org/), which contains 1443 verified position weight matrices from the HOCOMOCO H12CORE collection [75]. Motifs were searched for within the 500 bp upstream and 50 bp downstream regions of the *VLG-1* transcription start site. Matched motifs were considered as hits after multiple testing correction using the Bonferroni method. Before visualization of the motifs on the *VLG-1* promoter, redundancy was addressed by merging hits from the same TF family overlapping more than 50% in position, as well as merging similar TF families into categories.

### Statistics

Gene expression data were analyzed by one-way analysis of variance (ANOVA) after log_10_- transformation of the 2^-dCt^ values, followed by Tukey-Kramer’s Honest Significant Difference (HSD) test. Viral loads, knockdown efficiency, and fecundity estimates were compared pairwise with a Mann-Whitney’s non-parametric test. Proportions (midgut prevalence, carcass prevalence, carcass-to-head dissemination prevalence, transmission efficiency, fertility) were analyzed using a chi-squared non-parametric test. Survival assays were analyzed with a Gehan-Breslow-Wilcoxon test. Gene set enrichment analysis in the transcriptomic dataset was performed with Fisher’s statistical test. Only genes with FDR < 0.05 were considered significantly enriched. Statistical analyses were performed in Prism v.10.1.0 (www.graphpad.com), JMP v.14.0.0 (www.jmp.com), and R v.4.3.2 (www.r-project.org).

### Data availability

The data discussed in this publication have been deposited in NCBI’s Gene Expression Omnibus [76] and are accessible through GEO Series accession number GSE269945.

## ACKNOWLEDGEMENTS

We thank Catherine Lallemand for assistance with mosquito rearing, Artem Baidaliuk for preliminary bioinformatic analysis of *Vago*-like gene homology, and Prasad Paradkar for kindly sharing the CxVLG-1 antibody. RNA-seq library preparation was performed by Elodie Turc and Laure Lemée from the Biomics platform (C2RT, Institut Pasteur, Paris, France) supported by France Génomique (ANR-10-INBS-09) and IBISA. This work was supported by the French Government’s Investissement d’Avenir program, Laboratoire d’Excellence Integrative Biology of Emerging Infectious Diseases (grant ANR-10-LABX-62-IBEID to L.L. and E.C.), Agence Nationale de la Recherche (grant ANR-18-CE35-0003-01 to L.L.), a PhD grant from Ecole Normale Supérieure de Lyon (to E.C.) and a junior seed grant from Institut Pasteur (to E.C.). The funders had no role in study design, data collection and analysis, decision to publish, or preparation of the manuscript.

## AUTHOR CONTRIBUTIONS

Conceptualization: E.C., J.D., P.M., T.V., S.H.M., L.L.

Investigation: E.C., A.B.C., J.D., A.B., F.A.H.v.H., U.P., T.V., S.D.

Data curation: H.V.

Formal analysis: E.C., H.V., J.D., F.A.H.v.H., A.B., U.P., L.L.

Visualization: E.C., F.A.H.v.H., H.V.

Writing – original draft: E.C., S.H.M., L.L.

Writing – review and editing: E.C., P.M., S.H.M., L.L.

Funding acquisition: E.C., S.H.M., L.L.

## SUPPLEMENTARY FIGURES LEGENDS

**Supplementary figure S1.**
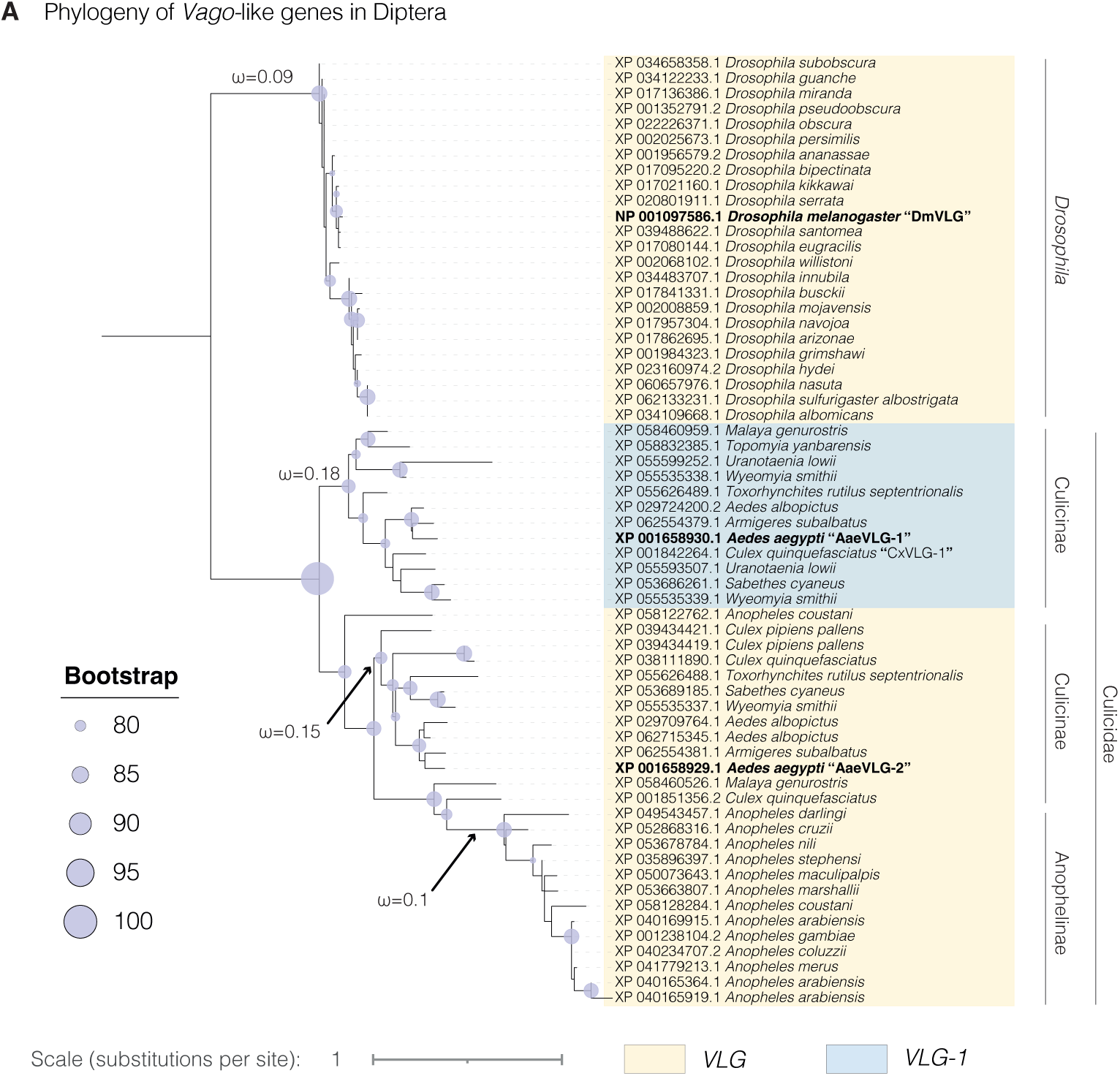
Phylogenetic tree of *Vago*-like gene homologs among Culicidae and *Drosophila* species. The tree was constructed with a maximum-likelihood analysis of amino-acid sequences with at least 30% identity with *D. melanogaster VLG* (*DmVLG*, CG14132). Accession number (RefSeq) of the homolog protein and name of the species are indicated at the tip of each branch. *VLG* and *VLG-1* clades are colored in yellow and blue, respectively. The size of blue dots represents the bootstrap support of each node. The dN/dS (ω) estimates are indicated for the main branches.

**Supplementary figure S2.**
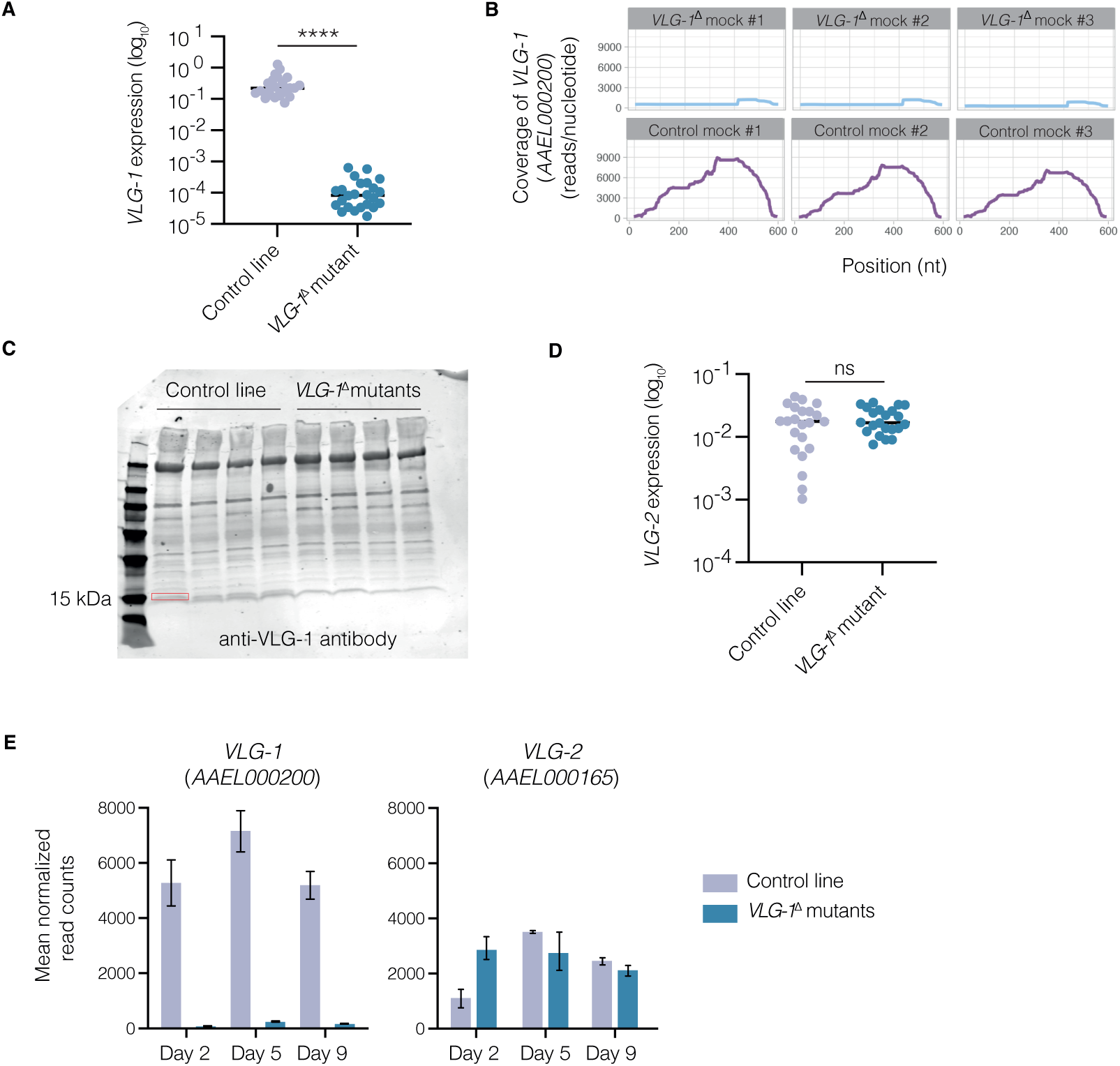
Evidence for *VLG-1* loss of function in mutant *Ae. aegypti*. (A) *AaeVLG-1* transcript expression levels detected by RT-qPCR in the control and *VLG-1*^Δ^ mutant lines. ****p<0.0001 (Mann-Whitney’s test) (B) Coverage (number of reads per nucleotide position) of *AaeVLG-1* transcripts by RNA-seq in *VLG-1*^Δ^ mutants (top panels) and controls (bottom panels) in three pools of 10 bodies for each line. (C) Western blotting of VLG-1 protein using an anti-CxVLG-1 antibody in controls and *VLG-1*^Δ^ mutants, in four pools of five females for each line. The band corresponding to VLG-1 theoretical size (15 kiloDaltons (kDa)) is highlighted in red and is detected in the controls but not in the *VLG-1*^Δ^ mutants. (D) *AaeVLG-2* transcript expression levels detected by RT-qPCR in the control and *VLG-1*^Δ^ mutant lines. (E) Number of reads of *AaeVLG-1* and *AaeVLG-2* transcripts detected by RNA-seq in bodies of controls and *VLG-1*^Δ^ mutants on days 2, 5, and 9 after a mock bloodmeal. Mean normalized counts (obtained with DESeq2 [71]) from three pools of 10 bodies for each line are depicted. Vertical bars represent standard deviations. In (A) and (D), gene expression levels are normalized to the transcript abundance of the housekeeping gene encoding ribosomal protein S 17 (*RPS17*), and expressed as 2^-dCt^, where dCt = Ct*_Gene_* – Ct*_RPS17_*.

**Supplementary figure S3.**
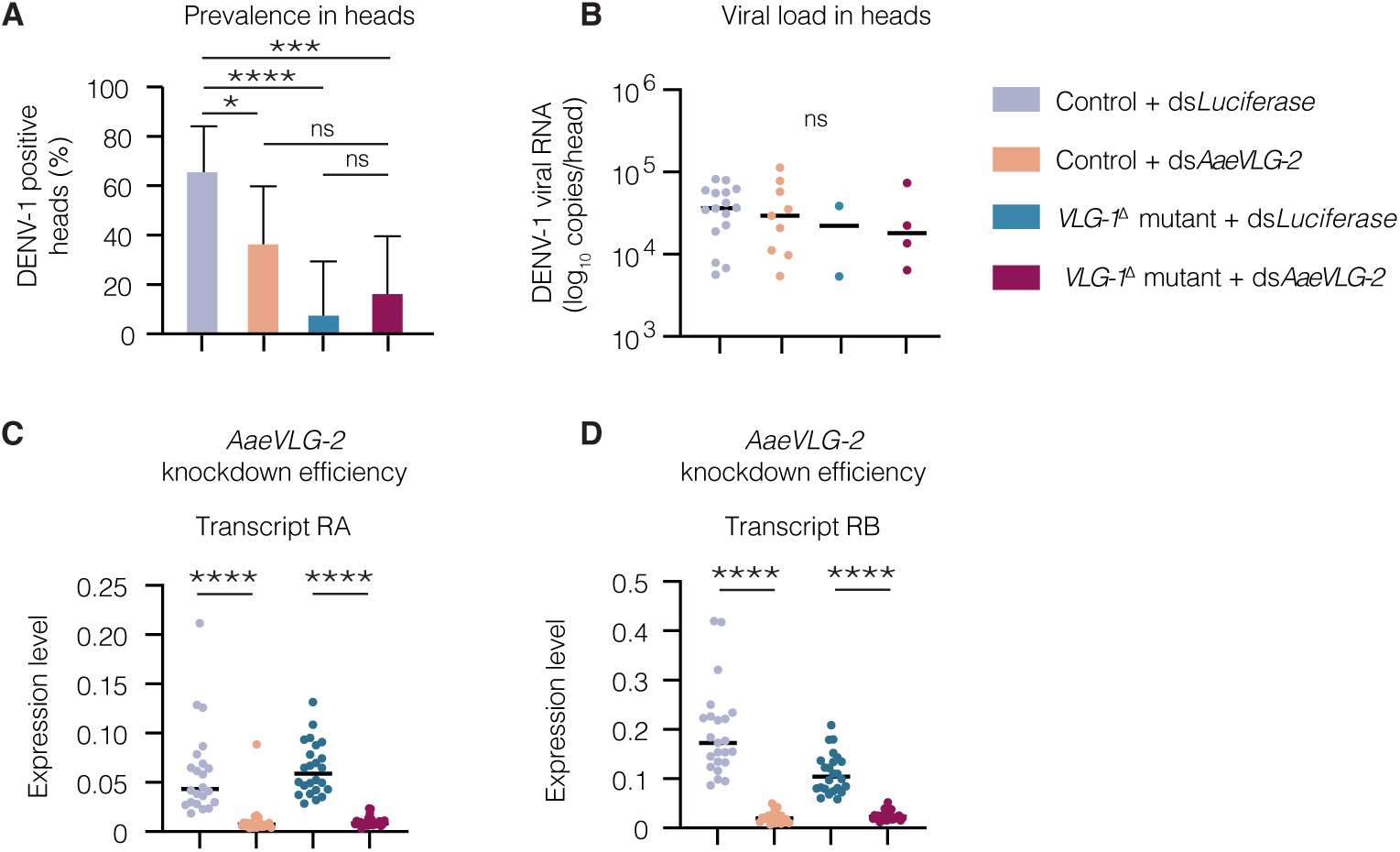
*AaeVLG-2* knockdown reduces the proportion of DENV-positive heads. Female *Ae. aegypti* from both the *VLG-1*^Δ^ mutant line and the control line were injected with either dsRNA targeting *AaeVLG-2* or a dsRNA targeting the *Luciferase* gene as a negative control. Forty-eight hours after injection (on the day of the infectious bloodmeal), mosquitoes were offered an infectious bloodmeal containing 5×10^6^ FFU/mL of DENV-1. Heads were collected on day 7 after the bloodmeal and processed for viral RNA quantification to evaluate infection prevalence (A) and viral loads (B). In parallel, whole unfed mosquitoes were collected on the day of the bloodmeal to quantify *AaeVLG-2* transcript RA (C) and transcript RB (D) abundance by RT-qPCR. Gene expression levels are normalized to the transcript abundance of the housekeeping gene encoding ribosomal protein S 17 (*RPS17*), and expressed as 2^-dCt^, where dCt = Ct*_Gene_* – Ct*_RPS17_*. *p<0.05; **p<0.01; ***p<0.001 (Mann-Whitney’s test for gene knockdown efficiency and viral loads, chi-squared test for prevalence).

**Supplementary figure S4.**
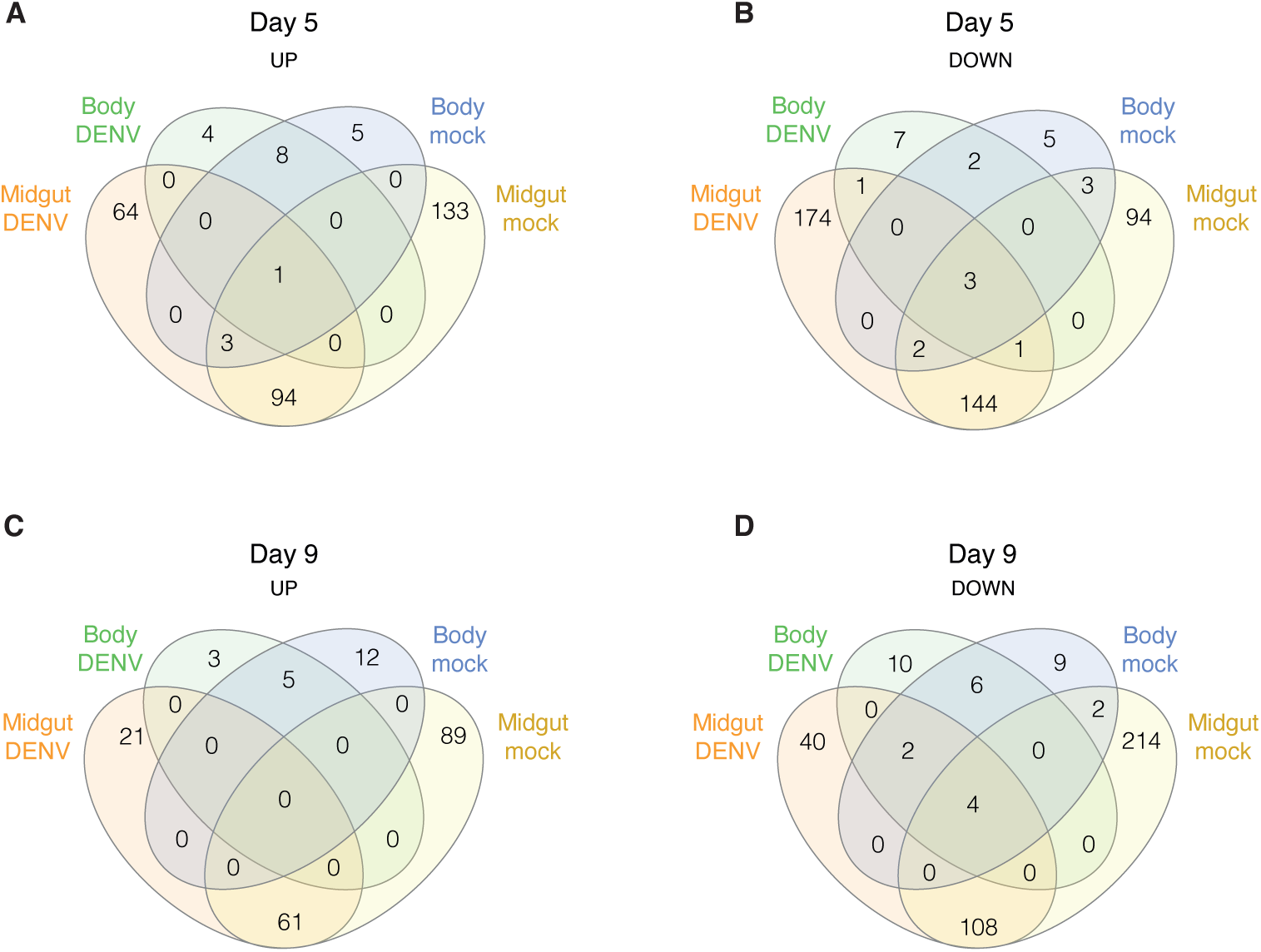
Overlap of differentially expressed genes in *VLG-1*^Δ^ mutants compared to wild-type controls on days 5 and 9 post bloodmeal. Venn diagrams show the number of up-regulated (A, C) and down-regulated (B, D) differentially expressed genes shared between experimental conditions on day 5 (A-B) and day 9 (C-D) post bloodmeal.

**Supplementary figure S5.**
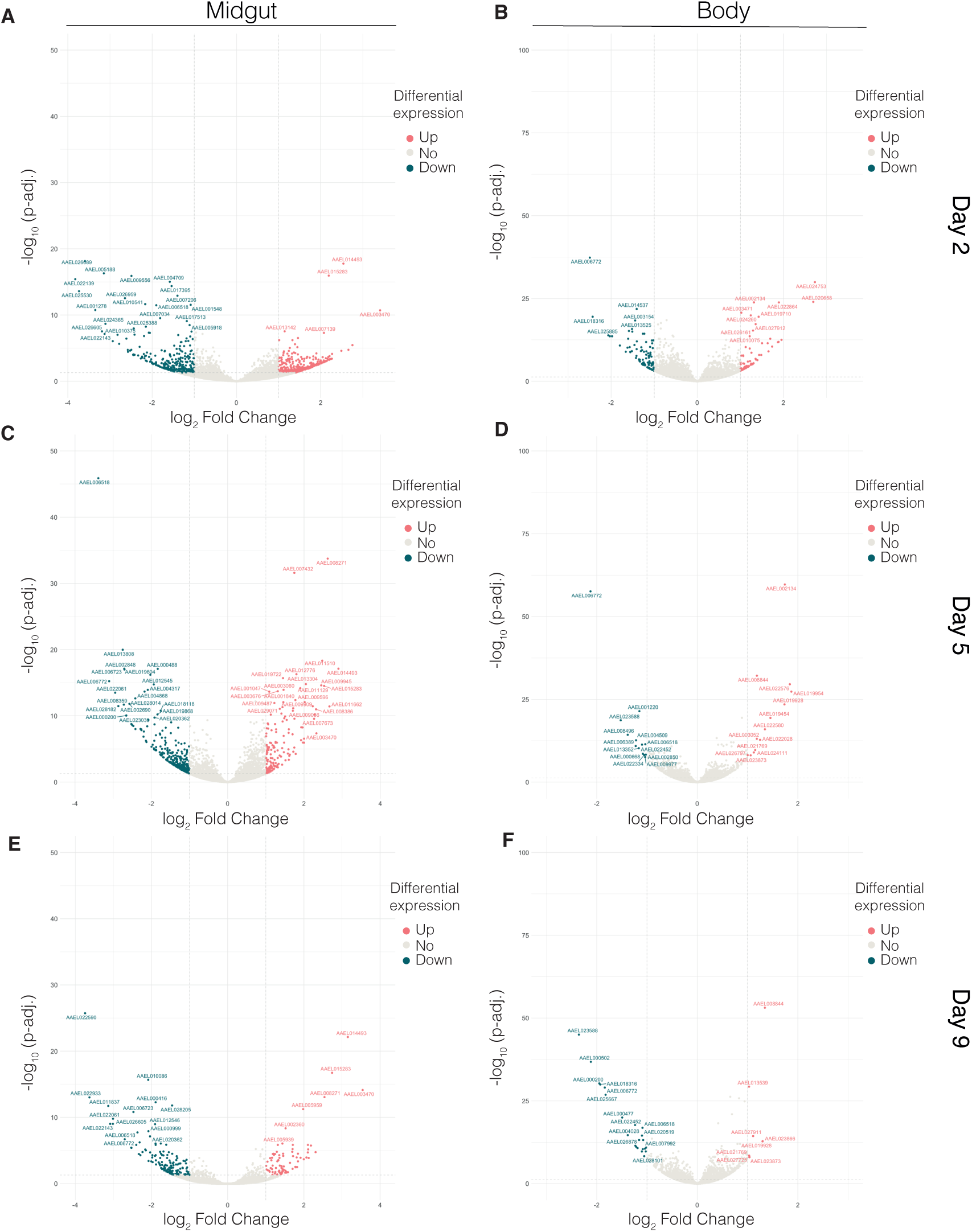
Volcano plots of differentially expressed genes in *VLG-1*^Δ^ mutants compared to wild-type controls in the DENV-exposed condition. Statistical significance of the difference in gene expression between mutants and controls (adjusted for multiple testing) is shown as a function of the log_2_-transformed fold change in expression. Genes that are significantly up-regulated and down-regulated are shown in red and blue, respectively. Comparisons were performed separately by tissue ((A, C, E): midgut; (B, D, F): body) and timepoint ((A-B): day 2; (C-D): day 5; (E-F): day 9 post bloodmeal). When detected, *AAEL000200* was removed from the plot to avoid graphical distortion due to its extremely low expression in *VLG-1*^Δ^ mutants.

**Supplementary figure S6.**
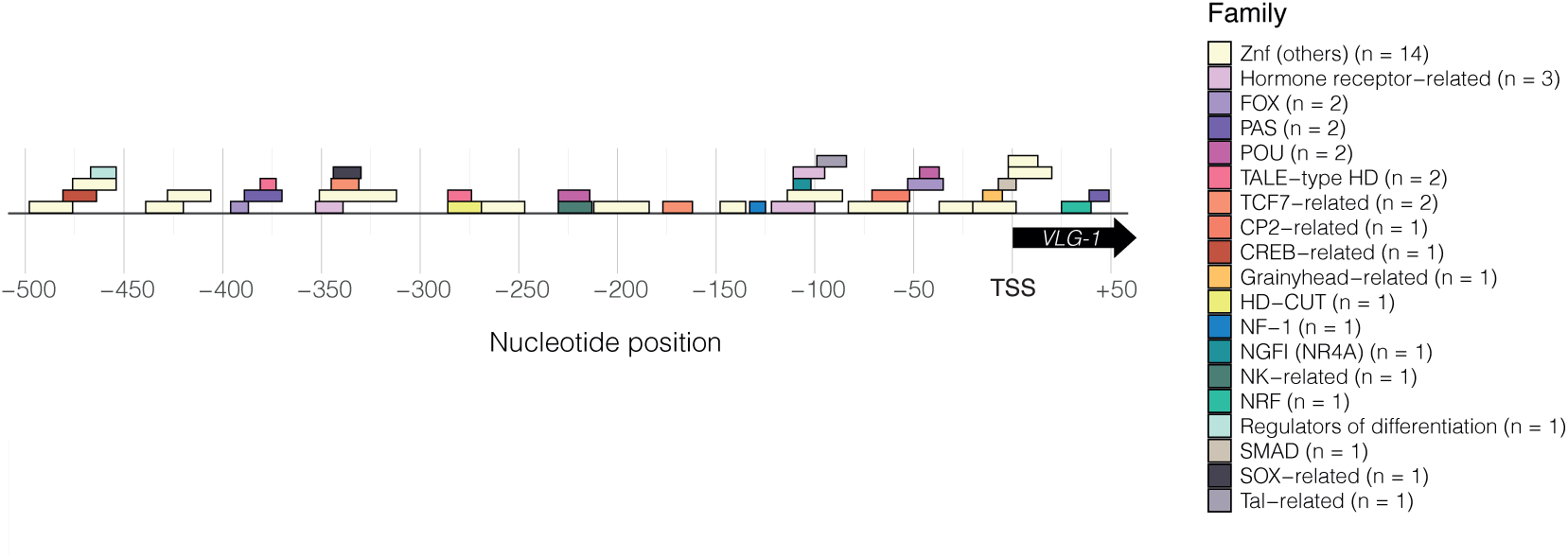
Transcription factor binding motifs found in the *Ae. aegypti VLG-1* promoter sequence. Motif hits identified and classified per transcription factor family in the promoter region (500 bp upstream and 50 bp downstream of the *VLG-1* transcription start site (TSS) (TSS coordinates: chr3:215,597,712). Nucleotide position is indicated relative to the TSS. The Znf (others) category includes the “Other with up to three adjacent zinc fingers”, “More than 3 adjacent zinc fingers” and “Multiple dispersed zinc fingers” transcription factor families. The “Hormone-receptor related” category includes the “Steroid hormone receptors”, “Thyroid hormone receptor-related” and “RXR-related receptors” families. The number of identified motifs is indicated for each motif category.

## SUPPLEMENTARY TABLES

**Supplementary Table 1.**
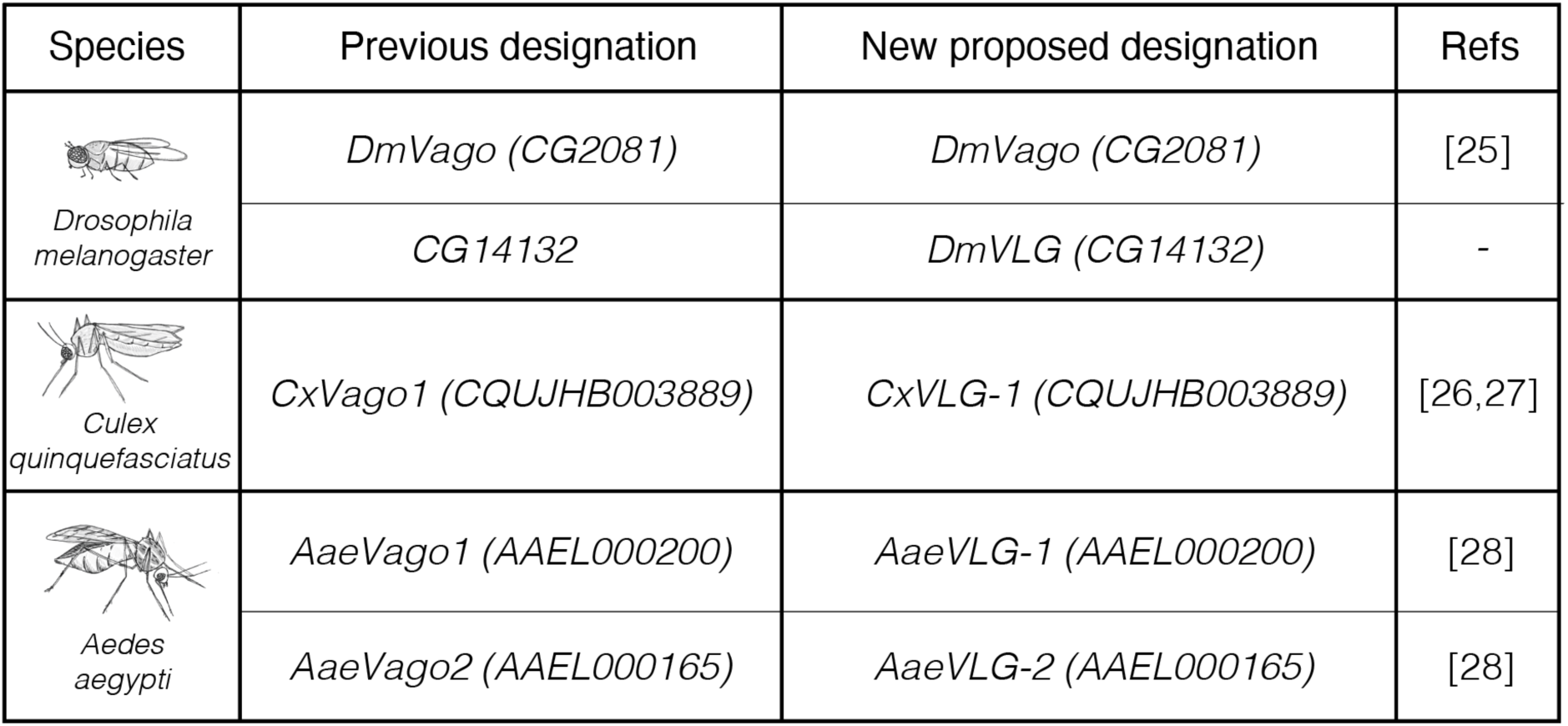
Proposed updated designation of *Vago* and *Vago*-like genes.

| Species | Previous designation | New proposed designation | Refs |
| --- | --- | --- | --- |
| *Drosophila melanogaster* | *DmVago* (CG2081) | *DmVago* (CG2081) | [25] |
|  | CG14132 | *DmVLG* (CG14132) | - |
| *Culex quinquefasciatus* | *CxVago1* (CQUJHB003889) | *CxVLG-1* (CQUJHB003889) | [26,27] |
| *Aedes aegypti* | *AaeVago1* (AAEL000200) | *AaeVLG-1* (AAEL000200) | [28] |
|  | *AaeVago2* (AAEL000165) | *AaeVLG-2* (AAEL000165) | [28] |

**Supplementary Table 2.**
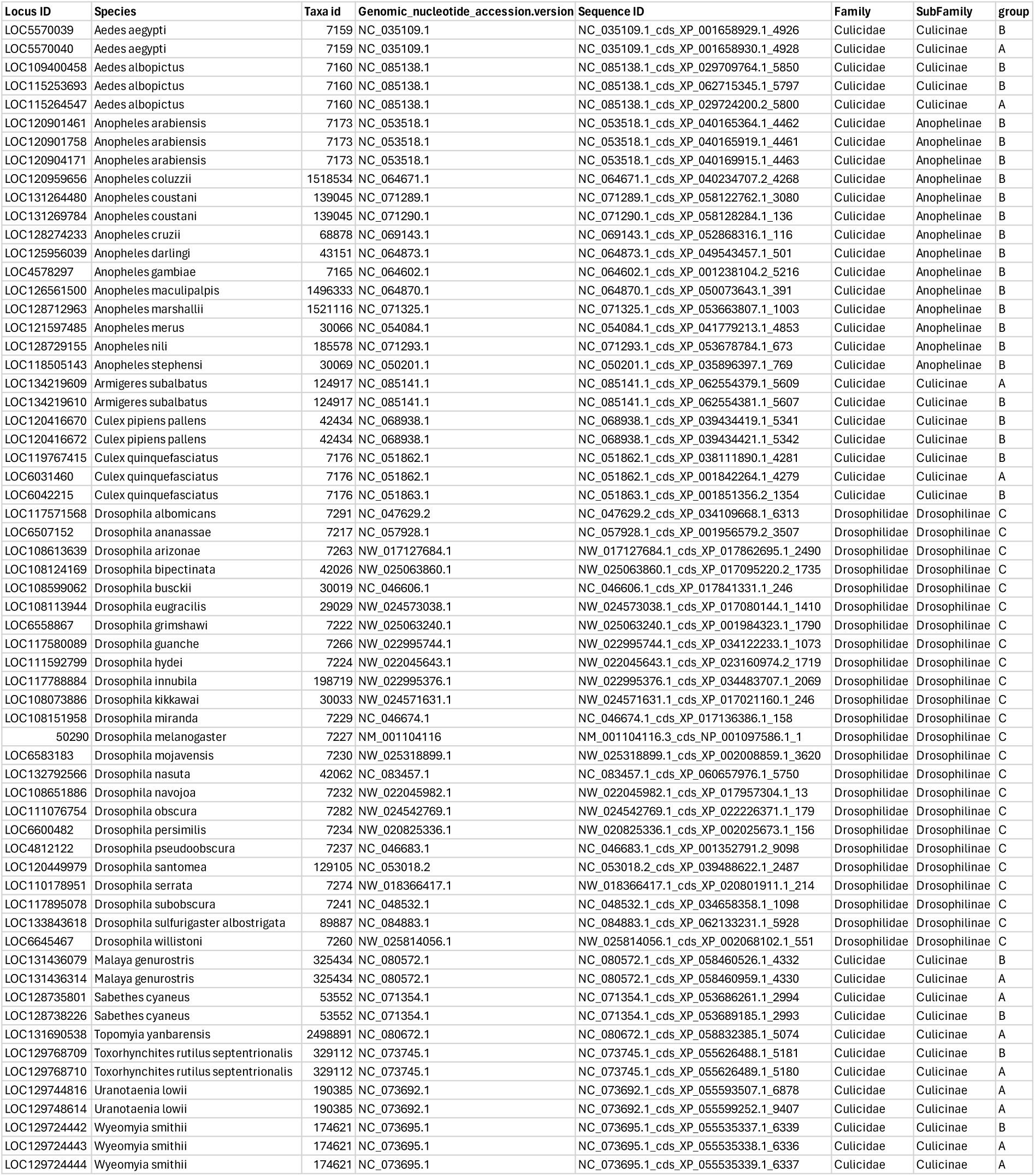
*Vago*-like gene homologs used in the gene phylogeny.

**Supplementary Table 3.** Results of dN/dS analysis in CODEML.

|  |  | outgroup | Vago1 | Vago (Culicinae) | Vago (Anophelinae) |  |  |  |  |
| --- | --- | --- | --- | --- | --- | --- | --- | --- | --- |
| model | likelihood | w0 | w1 | w2 | w3 | 2Δℓ | pvalue | df | model description |
| M0 | -7009,86928 | 0.12623 |  |  |  |  |  |  | M0: null model assumes the same w for all branches. |
| M1 | -7004,40804 | 0.106562 | 0.174765 |  |  | 10,922466 | 0.0009500506 | 1 | M1: foreground group is only Vago1 |
| M2 | -7002,76217 | 0.0853288 | 0.17507 | 0.120153 |  | 14,214216 | 0.0008192609 | 2 | M2: 2 foreground groups Vago1 and Vago Culicinae and Anophelinae |
| M3 | -7001,25308 | 0.0878059 | 0.175461 | 0.149161 | 0.0972423 | 17,232404 | 0.0006330647 | 3 | M3: 3 foreground groups (Vago1, Vago Culicinae and Vago Anophelinae) |
| # alpha critical value = 3.841459 |  |  |  |  |  |  |  |  |  |

**Supplementary Table 4.**
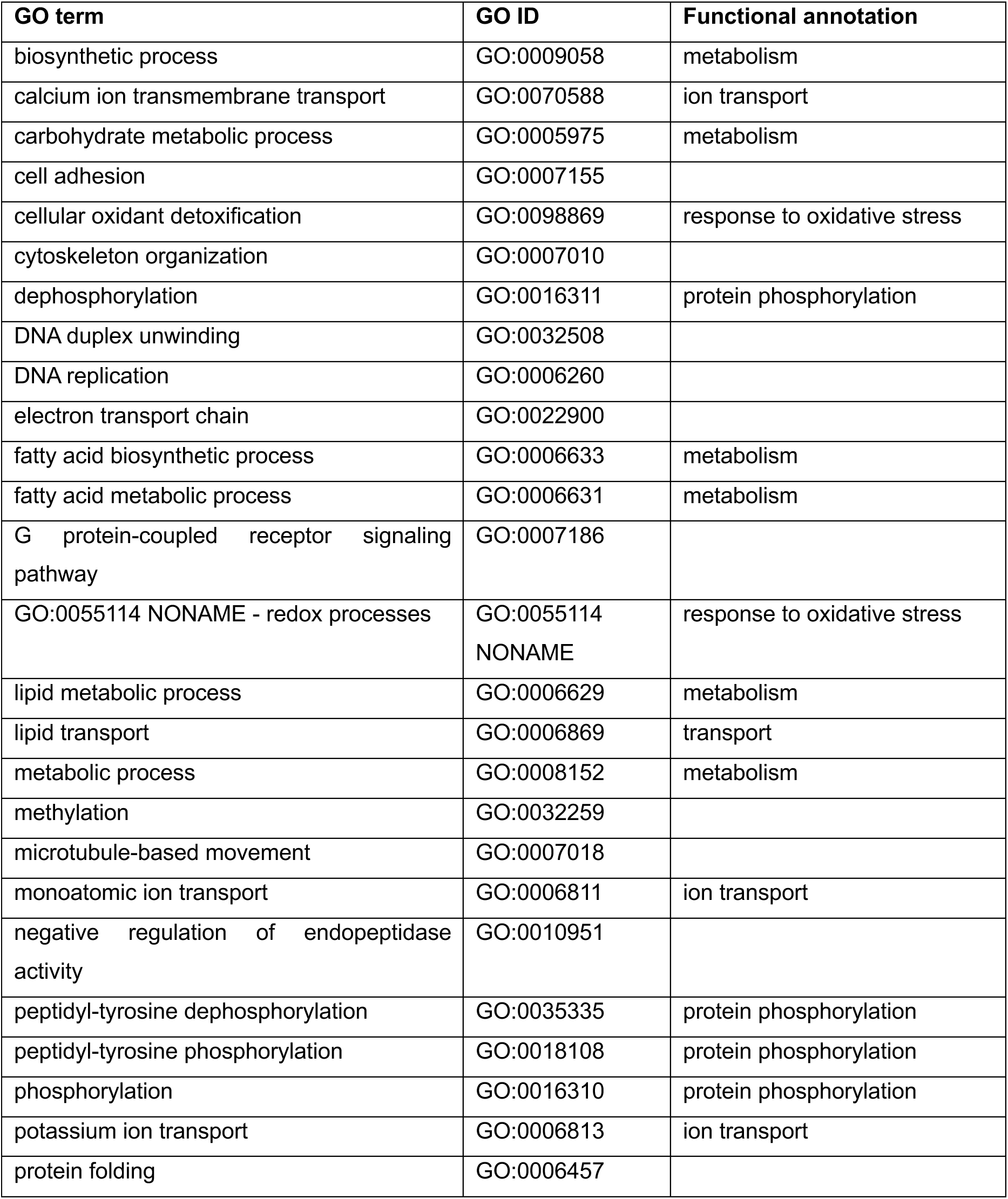
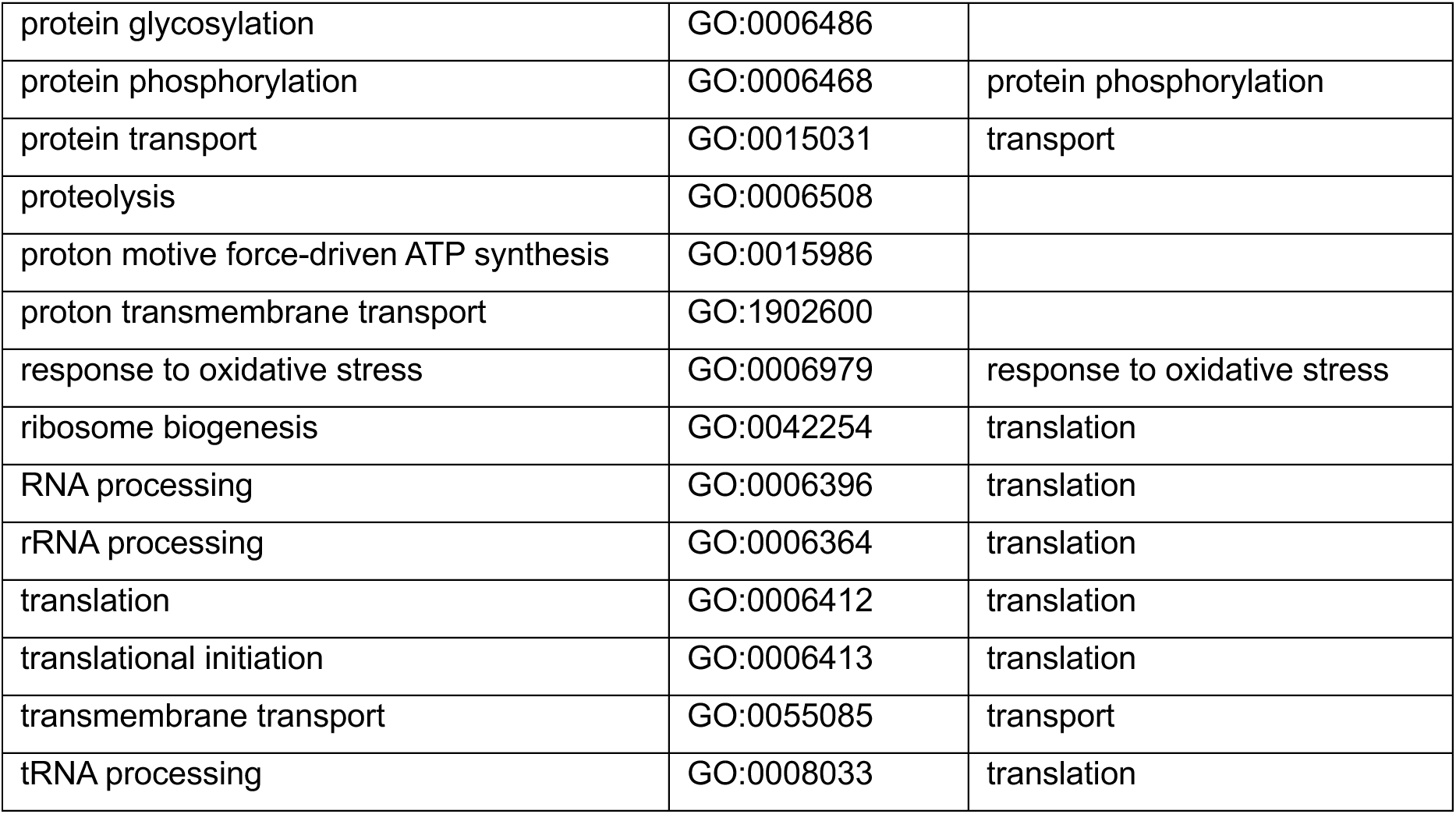
GO terms.

| GO term | GO ID | Functional annotation |
| --- | --- | --- |
| biosynthetic process | GO:0009058 | metabolism |
| calcium ion transmembrane transport | GO:0070588 | ion transport |
| carbohydrate metabolic process | GO:0005975 | metabolism |
| cell adhesion | GO:0007155 |  |
| cellular oxidant detoxification | GO:0098869 | response to oxidative stress |
| cytoskeleton organization | GO:0007010 |  |
| dephosphorylation | GO:0016311 | protein phosphorylation |
| DNA duplex unwinding | GO:0032508 |  |
| DNA replication | GO:0006260 |  |
| electron transport chain | GO:0022900 |  |
| fatty acid biosynthetic process | GO:0006633 | metabolism |
| fatty acid metabolic process | GO:0006631 | metabolism |
| G protein-coupled receptor signaling pathway | GO:0007186 |  |
| GO:0055114 NONAME - redox processes | GO:0055114<br>NONAME | response to oxidative stress |
| lipid metabolic process | GO:0006629 | metabolism |
| lipid transport | GO:0006869 | transport |
| metabolic process | GO:0008152 | metabolism |
| methylation | GO:0032259 |  |
| microtubule-based movement | GO:0007018 |  |
| monoatomic ion transport | GO:0006811 | ion transport |
| negative regulation of endopeptidase activity | GO:0010951 |  |
| peptidyl-tyrosine dephosphorylation | GO:0035335 | protein phosphorylation |
| peptidyl-tyrosine phosphorylation | GO:0018108 | protein phosphorylation |
| phosphorylation | GO:0016310 | protein phosphorylation |
| potassium ion transport | GO:0006813 | ion transport |
| protein folding | GO:0006457 |  |
| protein glycosylation | GO:0006486 |  |
| protein phosphorylation | GO:0006468 | protein phosphorylation |
| protein transport | GO:0015031 | transport |
| proteolysis | GO:0006508 |  |
| proton motive force-driven ATP synthesis | GO:0015986 |  |
| proton transmembrane transport | GO:1902600 |  |
| response to oxidative stress | GO:0006979 | response to oxidative stress |
| ribosome biogenesis | GO:0042254 | translation |
| RNA processing | GO:0006396 | translation |
| rRNA processing | GO:0006364 | translation |
| translation | GO:0006412 | translation |
| translational initiation | GO:0006413 | translation |
| transmembrane transport | GO:0055085 | transport |
| tRNA processing | GO:0008033 | translation |

**Supplementary Table 5.**
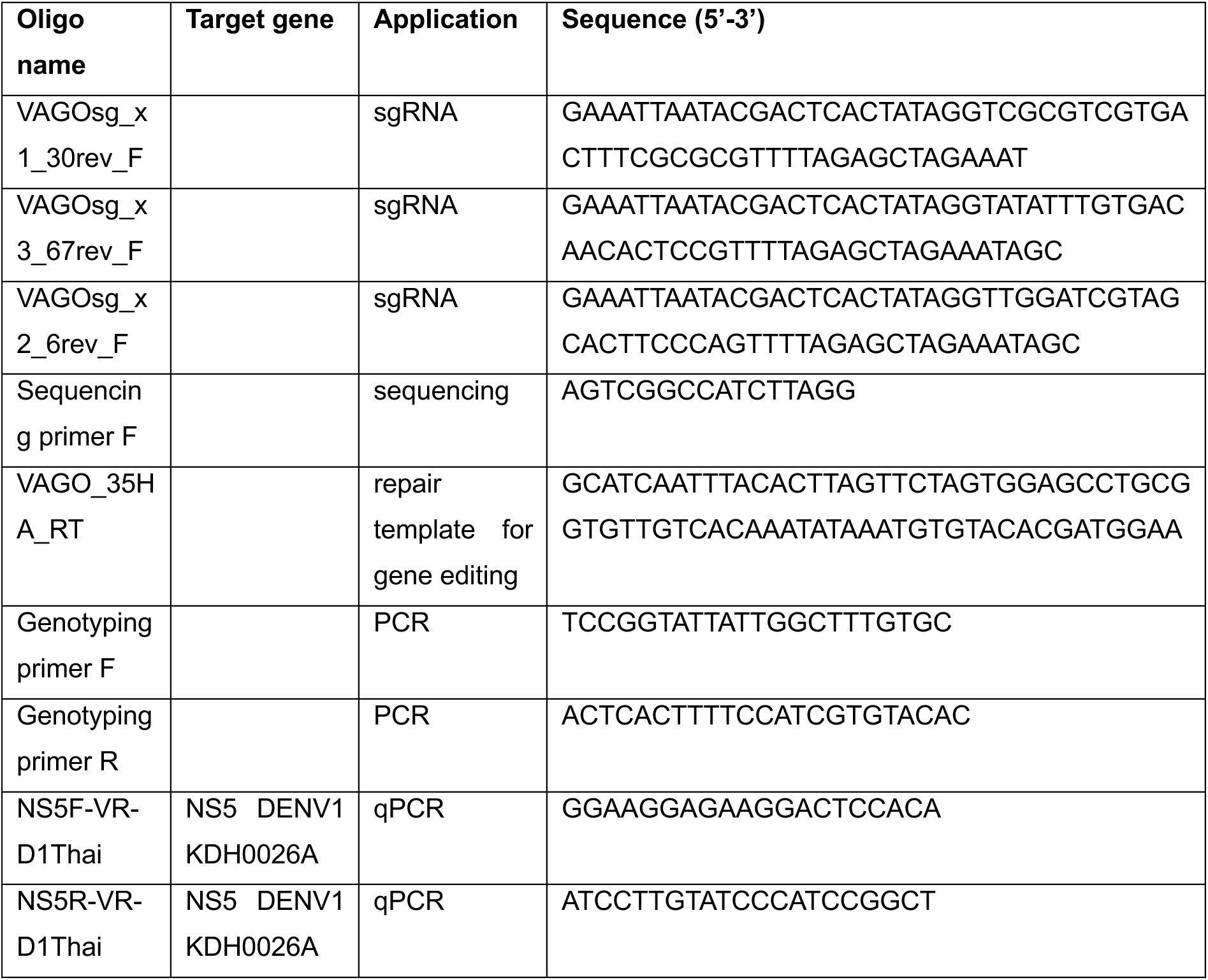
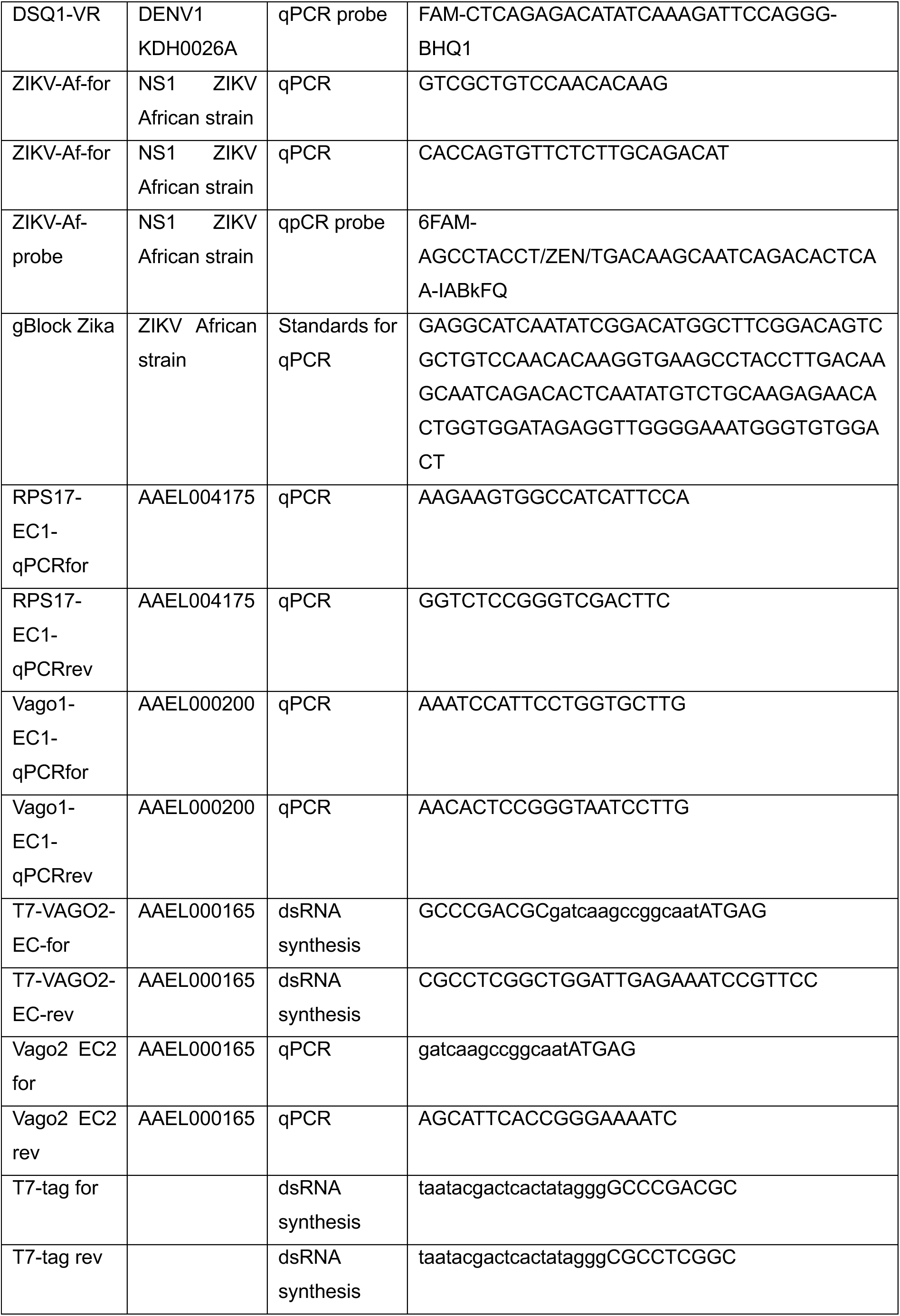
Oligonucleotide sequences.

**Supplementary Table 6:** Hits for transcription factor DNA binding motifs from HOCOMOCO H12CORE in the promoter of *VLG-1.* Start and end positions of motifs are indicated relative to *VLG-1* transcription start site.

| TF | Start | End | Strand | TF Family | p-value | Corrected p-value |
| --- | --- | --- | --- | --- | --- | --- |
| ZBTB49 | -498 | -476 | - | More than 3 adjacent zinc fingers | 3.311e-5 | 0.04778 |
| CREM | -481 | -464 | + | CREB-related | 1.312e-5 | 0.01893 |
| ZNF766 | -476 | -454 | - | More than 3 adjacent zinc fingers | 1.718e-5 | 0.02479 |
| MEF2B | -467 | -454 | + | Regulators of differentiation | 1.841e-5 | 0.02657 |
| ZNF615 | -439 | -420 | - | More than 3 adjacent zinc fingers | 1.510e-5 | 0.02179 |
| ZNF26 | -428 | -406 | - | More than 3 adjacent zinc fingers | 2.999e-5 | 0.04328 |
| FOXP3 | -396 | -387 | - | FOX | 2.624e-5 | 0.03786 |
| NPAS2 | -389 | -370 | + | PAS | 2.636e-5 | 0.03804 |
| IRX1 | -381 | -373 | - | TALE-type HD | 2.089e-5 | 0.03014 |
| CLOCK | -380 | -370 | - | PAS | 2.489e-5 | 0.03592 |
| NPAS2 | -380 | -370 | + | PAS | 2.636e-5 | 0.03804 |
| PGR | -353 | -339 | + | Steroid hormone receptors | 2.037e-5 | 0.02939 |
| ZIK1 | -351 | -329 | - | More than 3 adjacent zinc fingers | 3.155e-5 | 0.04553 |
| ZNF613 | -350 | -326 | + | More than 3 adjacent zinc fingers | 2.812e-6 | 0.00406 |
| ZNF570 | -348 | -330 | - | More than 3 adjacent zinc fingers | 1.268e-6 | 0.00183 |
| ZNF362 | -348 | -326 | - | More than 3 adjacent zinc fingers | 1.589e-5 | 0.02293 |
| LEF1 | -345 | -331 | + | TCF7-related | 1.274e-5 | 0.01838 |
| ZNF791 | -345 | -323 | + | More than 3 adjacent zinc fingers | 2.317e-5 | 0.03343 |
| SOX17 | -344 | -330 | + | SOX-related | 5.754e-6 | 0.00830 |
| ZNF362 | -343 | -321 | - | More than 3 adjacent zinc fingers | 1.995e-5 | 0.02879 |
| ZNF585A | -343 | -321 | - | More than 3 adjacent zinc fingers | 2.291e-5 | 0.03306 |
| ZNF716 | -342 | -318 | - | More than 3 adjacent zinc fingers | 2.612e-5 | 0.03769 |
| ZNF354A | -334 | -312 | + | More than 3 adjacent zinc fingers | 1.352e-5 | 0.01951 |
| ONECUT2 | -286 | -265 | - | HD-CUT | 2.163e-5 | 0.03121 |
| MEIS1 | -286 | -274 | - | TALE-type HD | 2.410e-5 | 0.03478 |
| ZNF432 | -269 | -247 | + | More than 3 adjacent zinc fingers | 3.013e-6 | 0.00435 |
| POU2F2 | -230 | -218 | - | POU | 9.311e-7 | 0.00134 |
| POU5F1B | -230 | -214 | + | POU | 2.084e-6 | 0.00301 |
| VENTX | -230 | -213 | - | NK-related | 2.118e-5 | 0.03056 |
| ZNF768 | -212 | -190 | + | More than 3 adjacent zinc fingers | 4.853e-6 | 0.00700 |
| ZNF490 | -206 | -184 | - | More than 3 adjacent zinc fingers | 2.443e-6 | 0.00353 |
| LEF1 | -177 | -163 | + | TCF7-related | 5.689e-6 | 0.00821 |
| TCF7L2 | -176 | -165 | + | TCF7-related | 1.161e-6 | 0.00168 |
| TCF7 | -176 | -162 | - | TCF7-related | 1.945e-6 | 0.00281 |
| TCF7L1 | -176 | -165 | + | TCF7-related | 2.296e-6 | 0.00331 |
| LEF1 | -176 | -165 | + | TCF7-related | 2.158e-5 | 0.03114 |
| YY2 | -148 | -135 | - | More than 3 adjacent zinc fingers | 4.667e-6 | 0.00673 |
| YY1 | -147 | -136 | - | More than 3 adjacent zinc fingers | 2.094e-5 | 0.03022 |
| NFIB | -133 | -125 | + | NF-1 | 3.027e-5 | 0.04368 |
| THRB | -122 | -101 | + | Thyroid hormone receptor-related | 2.203e-5 | 0.03179 |
| **ZNF558** | -114 | -90 | + | More than 3 adjacent zinc fingers | 2.692e-5 | 0.03885 |
| **NR2C2** | -111 | -100 | + | RXR-related receptors | 1.786e-5 | 0.02577 |
| **NR4A2** | -111 | -102 | + | NGFI (NR4A) | 2.051e-5 | 0.02960 |
| **NR2F2** | -111 | -100 | + | RXR-related receptors | 2.767e-5 | 0.03993 |
| **NR2F6** | -110 | -95 | + | RXR-related receptors | 1.368e-5 | 0.01974 |
| **NR2C1** | -110 | -95 | + | RXR-related receptors | 1.452e-5 | 0.02095 |
| **RARA** | -110 | -100 | + | Thyroid hormone receptor-related | 2.729e-5 | 0.03938 |
| **NR2C1** | -110 | -99 | + | RXR-related receptors | 3.342e-5 | 0.04823 |
| **TWIST1** | -99 | -84 | - | Tal-related | 5.000e-6 | 0.00722 |
| **MSC** | -98 | -87 | - | Tal-related | 2.312e-5 | 0.03336 |
| **TWIST2** | -96 | -87 | + | Tal-related | 5.224e-6 | 0.00754 |
| **ZBTB18** | -96 | -86 | + | More than 3 adjacent zinc fingers | 3.334e-5 | 0.04811 |
| **ZNF534** | -83 | -53 | - | More than 3 adjacent zinc fingers | 6.887e-6 | 0.00994 |
| **ZNF534** | -82 | -65 | - | More than 3 adjacent zinc fingers | 1.368e-5 | 0.01974 |
| **TFCP2L1** | -71 | -52 | + | CP2-related | 2.716e-5 | 0.03919 |
| **TFCP2L1** | -60 | -52 | - | CP2-related | 2.472e-5 | 0.03567 |
| **FOXB1** | -53 | -35 | + | FOX | 9.661e-6 | 0.01394 |
| **FOXC1** | -53 | -35 | + | FOX | 1.466e-5 | 0.02115 |
| **FOXA1** | -50 | -35 | + | FOX | 1.589e-5 | 0.02293 |
| **FOXA3** | -49 | -38 | + | FOX | 2.032e-5 | 0.02932 |
| **FOXA2** | -47 | -36 | + | FOX | 1.449e-5 | 0.02091 |
| **POU2F1** | -47 | -37 | - | POU | 1.538e-5 | 0.02219 |
| **POU5F1** | -47 | -37 | - | POU | 1.563e-5 | 0.02255 |
| **FOXA1** | -47 | -37 | + | FOX | 1.849e-5 | 0.02668 |
| **POU2F2** | -47 | -37 | - | POU | 2.228e-5 | 0.03215 |
| **FOXM1** | -47 | -36 | + | FOX | 2.710e-5 | 0.03911 |
| **ZSCAN1** | -37 | -20 | - | Other with up to three adjacent zinc fingers | 3.281e-5 | 0.04734 |
| **ZNF490** | -20 | 2 | - | More than 3 adjacent zinc fingers | 8.147e-6 | 0.01176 |
| **GRHL2** | -15 | -5 | - | Grainyhead-related | 9.661e-6 | 0.01394 |
| **SMAD2** | -7 | 2 | + | SMAD | 2.432e-5 | 0.03509 |
| **SMAD3** | -6 | 2 | - | SMAD | 2.858e-5 | 0.04124 |
| **ZNF787** | -2 | 13 | + | Multiple dispersed zinc fingers | 2.056e-5 | 0.02967 |
| **ZNF354A** | -2 | 20 | + | More than 3 adjacent zinc fingers | 2.228e-5 | 0.03215 |
| **NRF1** | 25 | 40 | - | NRF | 1.466e-5 | 0.02115 |
| **NPAS4** | 39 | 49 | - | PAS | 1.067e-6 | 0.00154 |

## Notes

### Competing Interest Statement

The authors have declared no competing interest.

### Summary of Updates

Title revised; No additional revisions in the text or the figures

